# Cortical microglia promote noradrenergic signaling to maintain wakefulness

**DOI:** 10.64898/2025.12.26.696634

**Authors:** Yue Liang, Wu Shi, Emily M. Dale, Shunyi Zhao, Fangfang Qi, Koichiro Haruwaka, Qixiu Hou, Mengdi Fei, Lauren Harris, Xiaoyang Hua, Long-Jun Wu

**Affiliations:** Center for Neuroimmunology and Glial Biology, Institute of Molecular Medicine, University of Texas Health Science Center at Houston, Houston, Texas, USA; Department of Neurology, Mayo Clinic, Rochester, Minnesota, USA; Mayo Clinic Graduate School of Biomedical Sciences, Rochester, Minnesota, USA; Department of Otorhinolaryngology, University of Texas Health Science Center at Houston, Houston, Texas, USA

## Abstract

Microglia are essential for maintaining brain homeostasis, including the sleep-wake cycle, but the underlying mechanisms remain unclear. Here, using in vivo two-photon imaging in both head-fixed and freely moving mice with simultaneous electroencephalogram–electromyogram (EEG–EMG) recording, we investigated microglial process dynamics and their interactions with locus coeruleus (LC) noradrenergic (NE) axons in the frontal cortex across sleep-wake states. During sleep or chemogenetic suppression of LC neurons, microglia enhanced their surveillance and increased interactions with NE axons. Correlative electron microscopy at nanometer resolution identified structural contacts between microglial process endings and NE boutons. Mechanistically, microglia-bouton interaction is primed by NE–β2-adrenergic receptor (β2AR) signaling, leading to enhanced Ca²⁺ activity in NE axons. These findings suggest that microglial interactions with NE axons promote cortical norepinephrine transmission and maintain stable wakefulness.

## Introduction

Sleep is a fundamental biological process that supports physiological homeostasis and overall health^1,2^. Over the past decades, significant progress has been made in mapping the neural circuitry underlying sleep–wake regulation, highlighting a highly coordinated network involving multiple brain regions and diverse neuronal subtypes^3,4^. While much of the current understanding has centered on neuronal mechanisms, accumulating evidence indicates that glial cells—particularly microglia—are emerging as active modulators of brain states under physiological conditions^5–7^.

Microglia, the resident immune cells of the central nervous system (CNS), have long been recognized for their roles in neurological disorders and in neurodevelopment^8–10^. Beyond these established functions, microglia maintain homeostasis with their dynamic surveillance behavior, continuously extending and retracting their processes to monitor and regulate the neural environment^11,12^. Moreover, microglial interactions with active neuronal elements often involve frequent and prolonged contacts, contributing to synaptic plasticity and local circuit synchronization^13–16^.

Given that sleep–wake stability is a fundamental aspect of brain homeostasis, it is crucial to understand how microglia might contribute to its regulation. Several studies have reported that microglial depletion—either pharmacologically or genetically—leads to a reduction in wake duration and an increase in Non-Rapid Eye Movement (NREM) sleep^17–19^, suggesting a role for microglia in maintaining sleep–wake balance. However, the dynamics of microglial processes and their functions across different brain states remain unclear. Microglia have been shown to influence the sleep–wake cycle by modulating thalamic reticular neuron excitability via ceramide signaling^19^ and by adapting CX3CR1 expression in response to the light/dark cycle to regulate synaptic activity^17^. In addition, increased microglial Ca²⁺ activity has been linked to sleep promotion^20^. Nevertheless, the mechanisms by which microglia regulate sleep–wake states are still not fully understood.

Our previous work together with others revealed that microglia enhance their surveillance and increase interactions with neuronal dendrites under anesthesia via NE–β2AR signaling^21,22^. Moreover, microglia extend their processes to shield axosomatic inhibitory inputs and transiently promote neuronal hyperactivity during emergence, a process that is dependent on NE–β2AR-mediated microglial process extension^23^. Considering that the central role of NE as a neuromodulator of arousal^24,25^ and as a regulator of neuron–microglia interactions^21–23^, we hypothesized that microglia modulate NE transmission, thereby contributing to sleep–wake regulation. To this end, we employed in vivo two-photon imaging in both head-fixed and freely moving mice, coupled with simultaneous EEG-EMG monitoring, to track microglial dynamics and their interactions with NE axons in the frontal cortex across sleep and wake states. We found that microglia increase their contacts with NE axon boutons through the NE–β2AR signaling during sleep, and that these interactions enhance Ca²⁺ activity in NE axons to maintain arousal stability.

## Result

### Microglia increase process surveillance during sleep

Previous studies have shown that microglia are important for modulating sleep–wake cycle^17,18^. Consistently, we found that mice treated with PLX3397 chow, which effectively depletes microglia (Figures S1A–C), showed a decreased amount of wakefulness and an increased NREM sleep during the dark phase (Figures S1D–F). These mice also exhibited shorter wake bout durations, accompanied by an increased number of both wake and NREM sleep bouts, as well as more frequent transitions between wakefulness and NREM sleep (Figures S1G–J). These findings indicate that microglial depletion disrupts the normal sleep–wake cycle by reducing wakefulness.

Microglial process dynamics, characterized by the continuous extension and retraction of their ramified processes, play a critical role in maintaining CNS homeostasis^26–28^. To investigate microglial process dynamics across sleep and wake states, we performed in vivo two-photon imaging in *Cx3cr1*^GFP/+^ mice in which microglia are labeled with GFP^29^. A chronic cranial window was implanted over the frontal cortex, and simultaneous EEG-EMG recording was established to classify behavioral states (Figure 1A). Ten-minute imaging epochs were selected based on vigilance state classification: periods with >80% wakefulness were defined as the wake phase, and those with >80% NREM sleep were defined as the sleep phase (Figure 1B, C). We found that during the sleep phase, microglia exhibited significantly increased surveillance territory (Figure 1D, E and Video S1) and occupation area (Figure 1F, G) compared to the wake phase. To rule out potential artifacts caused by head restraint and to capture continuous vigilance states, we next employed a miniaturized two-photon microscopy in freely-moving mice (Figure 1H). Similarly, microglia in freely-moving animals displayed enhanced surveillance territory and larger occupation area during sleep compared to wakefulness (Figure 1I-M). Together, these results demonstrate that microglial process dynamics are dependent on the vigilance states and are significantly enhanced during sleep.

**Figure 1.**
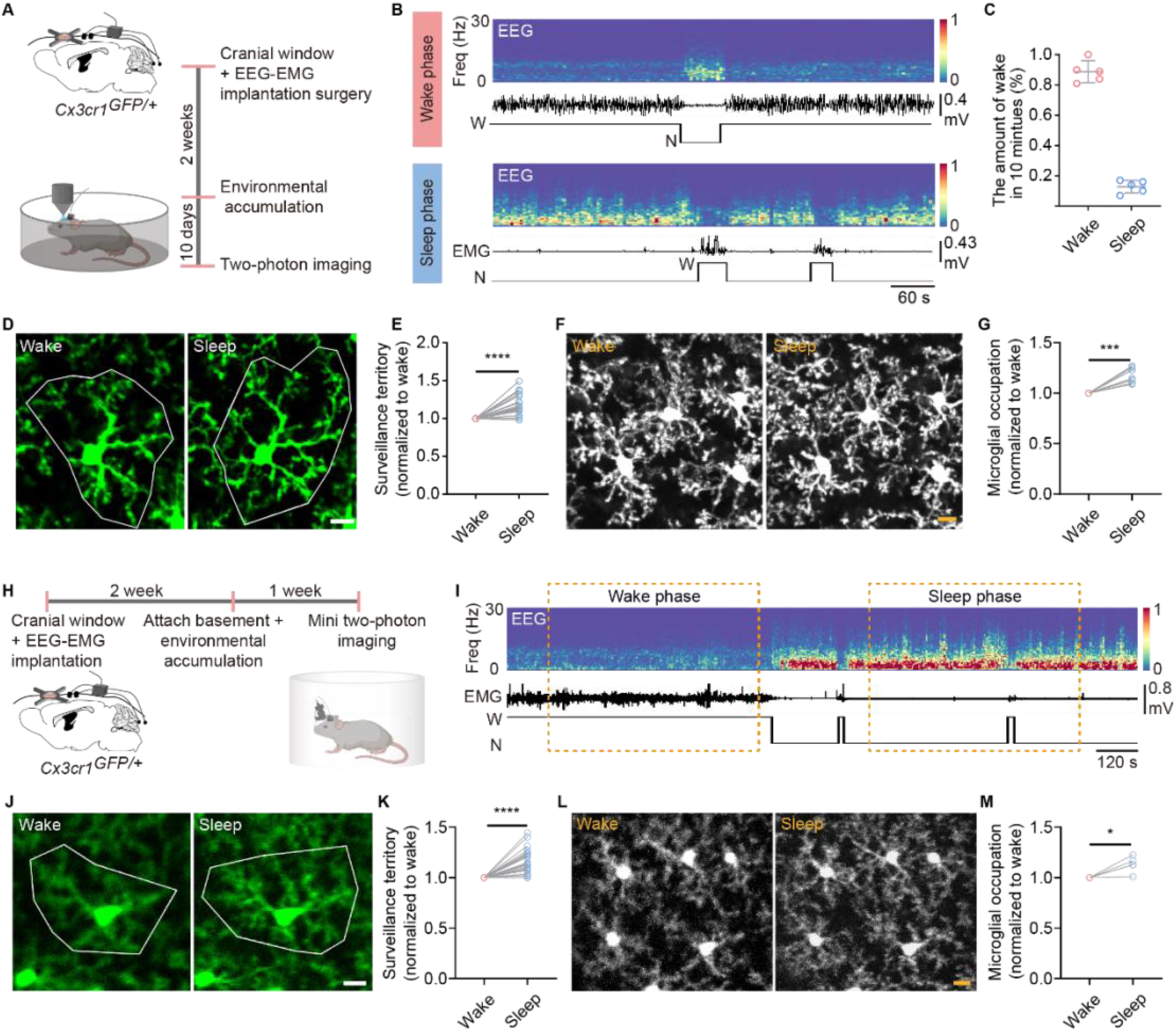
Microglia exhibit enhanced surveillance during sleep. (A) Schematic of the experimental procedure. (B) Representative EEG power spectrum, EMG trace, and hypnogram from one mouse during identified 10-minute wake and sleep periods. (C) The amount of wakefulness during the defined 10 minutes of wake and sleep (n=5 mice, data are presented as mean ± SD). (D) Representative two-photon images of microglia (GFP) during wake and sleep periods. The white line outline indicates the surveillance territory of the microglial process over a 10-minute period during wake and sleep states. Scale bars, 10 μm. (E) Quantification of microglial surveillance territory change during 10-minute wake and sleep states (n=23 microglia from 5 mice). (F) Representative two-photon images of microglia (max intensity projection) during 10-minute wake and sleep periods. Scale bars, 10 μm. (G) Quantification of microglial occupied area during 10-minute wake and sleep periods (n=5 mice). (H) Schematic of the experimental procedure for mini two-photon imaging. (I) Representative 1-hour EEG power spectrum, EMG trace, and hypnogram from a single mouse undergoing simultaneous mini two-photon imaging and EEG–EMG recording. The yellow box indicates the selected wake and sleep period used for imaging data analysis.9 (J) Representative mini two-photon images of microglia (GFP) during wake and sleep periods. The white line outline indicates the surveillance territory of the microglial process over a 10-minute period during wake and sleep states. Scale bars, 10 μm. (K) Quantification of microglial surveillance territory change during 10-minute wake and sleep states (n=30 microglia from 5 mice). (L) Representative mini two-photon images of microglia (max intensity projection) during 10-minute wake and sleep periods. Scale bars, 10 μm. (M) Quantification of microglial occupied area during 10-minute wake and sleep periods (n=5 mice). Data are presented as mean ± SEM, except for (C). Statistical significance was determined using a paired t-test in all graphs, *P < 0.05; **P < 0.01; *** P < 0.001; **** P < 0.0001. Each circle indicates an individual mouse.

### Enhanced microglial interactions with NE axons during sleep

NE plays a key role in regulating wakefulness^25,30^. To monitor NE level during the sleep-wake cycle, we virally expressed NE biosensor (AAV2/9-hSyn-GRAB-NE2h) in cortical neurons and then employed in vivo two-photon imaging in the frontal cortex. Indeed, we observed that NE levels display brain-state-dependent fluctuations, decreasing progressively from wakefulness to NREM, and further to REM sleep (Figure S2B–D). Consistently, fiber photometry recording also showed that NE level increased during wakefulness and decreased further during NREM and REM sleep in free-moving mice (Figure S2E-G).

The locus coeruleus (LC) is the primary source of cortical NE^31–33^ that influences microglial surveillance^21,22^. Thus, we investigated whether microglial interactions with LC-NE axons differ between sleep and wake states. To visualize LC-NE axons in the frontal cortex, we injected a mixture of AA9-TH-Cre and AAV9-CAG-FLEX-tdTomato into *Cx3cr1^GFP/+^* mice, allowing selective labeling of LC-NE axons with tdTomato. Immunohistochemical analysis confirmed the specificity of this labeling strategy: 88% of tdTomato-labeled axons were positive for dopamine beta-hydroxylase (DBH), and approximately 70% of NE axons were tdTomato-positive in the frontal cortex (Figure S3). Using in vivo two-photon imaging, we simultaneously visualized microglia (labeled with GFP) and NE axons (labeled with tdTomato) in the frontal cortex across sleep-wake cycles (Figure 2A). Average intensity projection images were generated for both wake and sleep phases (10 min each) to visualize microglial-NE axon interactions (Figure 2B). To characterize these interactions, we categorized microglial contacts with NE axons into three types: contacts with the microglial soma, with microglial branches, and with microglial process endings (Figure 2C). Quantitative analysis revealed a significant increase in contacts between NE axons and both microglial branches and process endings during the sleep phase (Figure 2D). To further assess the extent of these interactions, we manually segmented NE axons and quantified GFP fluorescence intensity along the segments in both wake and sleep phases (Figure 2E). The data showed that NE segments located in the border of microglial surveillance territories (segments in red) exhibited higher levels of GFP coverage during sleep (Figure 2F-H), suggesting enhanced microglial engagement.

**Figure 2.**
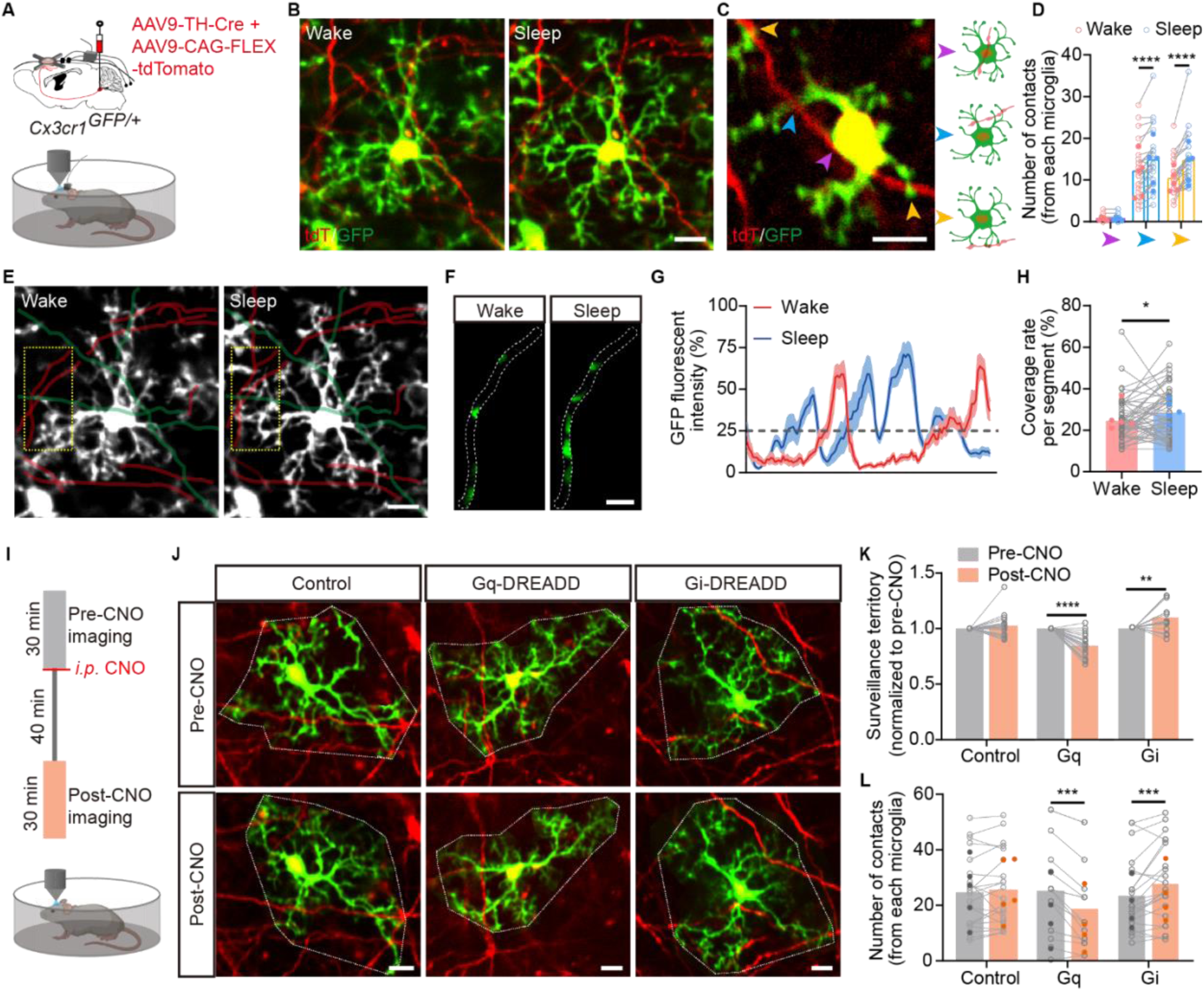
Enhanced microglial interactions with NE axons during sleep or in response to decreased norepinephrine levels. (A) Schematic of the experimental design for examining microglial interactions with NE axons during wake and sleep periods. (B) Representative two-photon images (average intensity projection of 10-minute imaging) showing microglia (GFP) and NE axons (tdTomato) during wake and sleep periods. Scale bar, 10 μm. (C) Representative image illustrating three types of microglial interactions with NE axons: microglial soma contacting NE axons (purple arrowhead), microglial branches contacting NE axons (blue arrowhead), and microglial process endings contacting NE axons (yellow arrowhead). Scale bar, 5 μm. (D) Quantification of the number of contacts between each microglia and NE axons for different contact types during wake and sleep periods (n=24 microglia from 5 mice). (E) Representative image showing microglia and NE axon segments during wake and sleep periods. Green NE segments are located within the central surveillance area of microglia, whereas red NE segments are located at the boundary. Yellow boxes indicate NE segments shown in (F). Scale bar, 10 μm. (F) Representative GFP (microglia) fluorescence within an NE axon segment [boxed region in (E)] during wake and sleep periods. Scale bar, 5 μm. (G) Plots of GFP (microglia) fluorescence intensity in the NE segment are shown in (F) during wake and sleep periods. (H) Quantification of microglial coverage of NE segments located at the boundary of the surveillance territory (red segments in (E)) during wake and sleep periods (n = 55 segments from 5 mice). (I) Schematic of the experimental procedure for CNO injection and two-photon imaging. (J) Representative two-photon images (average intensity projection of 10-minute imaging) of microglia (GFP) and NE axons (tdTomato) before and after CNO injection in the control, Gq-DREADD, and Gi-DREADD groups. Scale bar, 10 μm. (K) Quantification of changes in microglial surveillance territory over 10 minutes in the three groups (Control, n = 20 microglia from 5 mice; Gq, n = 21 microglia from 4 mice; Gi, n = 22 microglia from 4 mice). (L) Quantification of microglial process-ending contacts with NE axons during 10 minutes in the three groups. (Control, n = 20 microglia from 4 mice; Gq, n = 16 microglia from 4 mice; Gi, n = 22 microglia from 4 mice). Data are presented as mean ± SEM. Statistical significance was determined using a paired t-test in all graphs, *P < 0.05; **P < 0.01; *** P < 0.001; **** P < 0.0001. Each circle indicates an individual microglia. Each point indicates an individual mouse in D, H, and L.

LC-NE neuronal activity is critical in regulating sleep-wake states^34,35^. Next, we employed chemogenetic approaches to selectively manipulate LC-NE neurons and assess responses of cortical microglia. To this end, AAV/9-TH-Cre virus is bilaterally injected into the LC region in either *Cx3cr1^GFP/+^*(control)*, Cx3cr1^GFP/+^: R26^hM3Dq/+^* (Gq-DREADD) or *Cx3cr1^GFP/+^: R26^hM4Di/+^* (Gi-DREADD) mice (Figure S4A). The efficacy of neuronal modulation by chemogenetic approaches was validated by c-Fos immunostaining in the LC. As expected, following clozapine N-oxide (CNO) administration, the Gq-DREADD group exhibited markedly higher numbers of c-Fos–positive NE neurons in the LC region, whereas Gi-DREADD activation reduced c-Fos expression compared to control (Figure S4B, C). Correspondingly, fiber photometry recordings and head-fixed two-photon imaging in the frontal cortex confirmed NE level modulation: NE level increased in Gq-DREADD mice and decreased in Gi-DREADD mice following CNO injection (Figure S4D-J). These manipulations resulted in altered microglial morphology as well: microglial complexity was significantly reduced in Gq-DREADD mice after CNO injection, whereas it increased in the Gi-DREADD mice (Figure S4K-M). In addition, we found increased colocalization (both area and puncta) of IBA1 (microglia) and DBH (NE axons) in the Gi-DREADD group, while reduced colocalization in Gq-DREADD mice compared to controls (Figure S4N-S).

To examine the real-time response of microglia to chemogenetic manipulations of LC-NE neurons, we used two-photon in vivo imaging of GFP-labeled microglia and tdTomato-labeled LC-NE axons (Figure 2I). We found that microglia surveillance territory was decreased in Gq-DREADD mice but increased in Gi-DREADD mice post-CNO administration (Figure 2J, K). Consistently, the number of microglial process-ending contacts with NE axons was reduced in Gq-DREADD mice but significantly elevated in Gi-DREADD mice (Figure 2J, L). Together, these findings demonstrate that reduced NE levels promote microglial process extension and enhance interactions with LC-NE axons, whereas elevated NE suppresses microglial surveillance and contact.

### Microglial processes make more contacts with NE axon boutons than with shafts

NE boutons at the axons serve as sites for NE release and reuptake. Given our findings that microglia exhibit increased interactions with NE axons during sleep, we next investigated whether microglial process–bouton interactions may vary across different brain states. To this end, NE axons were classified into axon shafts (non-varicose segments) and axon boutons (varicose structures) under two-photon in vivo imaging (Figure 3A). We quantified the time-lapse imaging of microglial process-ending contacts with each axon compartment during 10-minute epochs of wakefulness and sleep. Our data revealed that microglial processes made significantly more contacts with axon boutons during sleep compared to wakefulness, while the number of contacts with axon shafts remained unchanged (Figure 3B-D and Video S2). However, the average duration of these contacts did not differ significantly between 10-minute sleep and wake states for either boutons or shafts (Figure 3E). Microglial bulbous endings represent specialized structures that mediate microglia–neuron interactions^23,36^. Notably, we observed that more than 70% of the microglial processes contacting NE axons during both wakefulness and sleep were bulbous endings (Figure 3F, G). To directly assess how NE affects microglia–bouton interactions, we examined contacts following chemogenetic manipulation of LC-NE neurons. In Gi-DREADD mice, which displayed enhanced microglial surveillance after CNO treatment, microglial contacts with NE axons increased specifically at axon boutons, with no significant changes at axon shafts (Figure 3H-K) or in contact duration across groups (Figure 3L). Conversely, Gq-DREADD mice showed a reduction in microglia–bouton contacts (Figure 3N–Q), again without changes in contact duration after CNO treatment (Figure 3R). Notably, microglia–axon interactions were analyzed within a 10-minute time window, therefore, contact durations were constrained to this period and are likely underestimated. Furthermore, time-lapse imaging (30 minutes before and after CNO injection) revealed no bouton formation or elimination associated with microglial contacts in either Gi-DREADD or Gq-DREADD mice (Figure 3H, I, N, O). Consistently, in both groups, microglial bulbous endings exhibited a higher proportion of contacts with NE boutons than non-bulbous endings, both before and after CNO administration (Figure M, S). Together, these results indicate that during extended microglial surveillance—either occurring during sleep or induced by chemogenetic inhibition of NE tone—microglial bulbous endings preferentially increase contacts with NE axon boutons.

**Figure 3.**
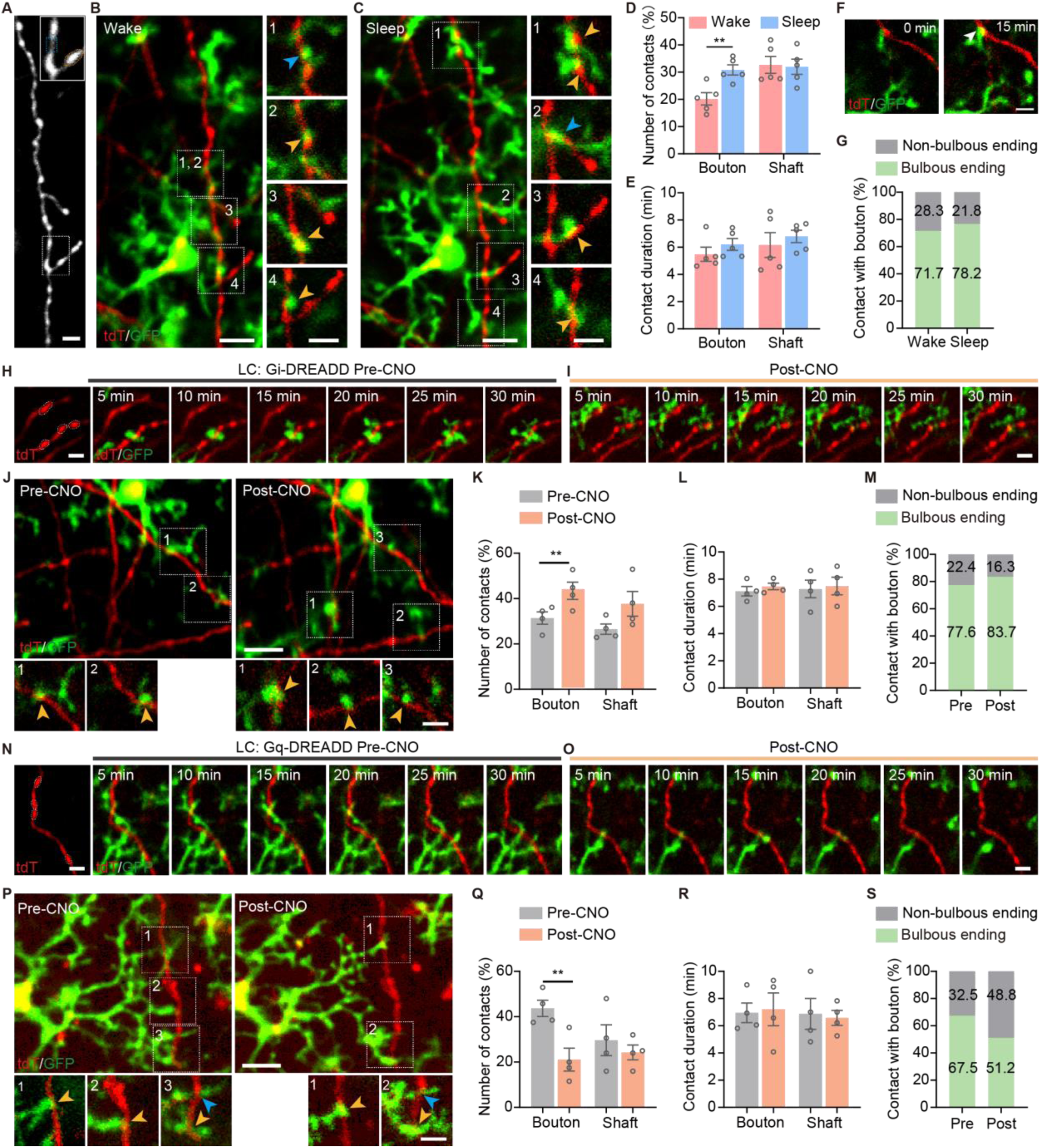
Microglial processes make more contact with NE axonal boutons than with axonal shafts. (A) Representative two-photon image of NE axons labeled with tdTomato. The white box indicates the region shown at higher magnification in the upper right corner. Yellow circles mark axonal boutons; blue boxes mark axonal shafts. Scale bar, 5 μm; magnified images, 5 μm. (B, C) Representative two-photon images of microglia (GFP) interacting with NE axons (tdTomato) during wake (B) and sleep (C) periods. White boxes indicate magnified regions in the right panels. Blue arrowheads mark microglial process endings in contact with NE axonal shafts; yellow arrowheads mark contacts with boutons. Scale bar, 10 μm; magnified images, 5 μm. (D, E) Quantification of the number (D) and duration (E) of microglial process endings in contact with NE axonal boutons or shafts during wake and sleep (n = 5 mice). (F) Representative time-lapse images acquired under a normal sleep–wake cycle showing microglia (GFP) interacting with NE axons (tdTomato). Arrowheads indicate microglial bulbous endings contacting NE boutons. Scale bar, 5 μm. (G) Proportion of non-bulbous and bulbous microglial endings contacting NE boutons during wakefulness and sleep (n = 5 mice). (H, I) Representative time-lapse images of microglia (GFP) interacting with NE axons (tdTomato) before (F) and after (G) CNO injection in the Gi-DREADD group. Scale bar, 5 μm. (J) Representative two-photon images of microglia (GFP) interacting with NE axons (tdTomato) before (left) and after (right) CNO injection in the Gi-DREADD group. White boxes indicate magnified regions at the bottom of panels; yellow arrowheads mark bouton contacts. Scale bar, 10 μm; magnified images, 5 μm. (K, L) Quantification of the number (K) and duration (L) of microglial process contacts with NE axonal boutons or shafts before and after CNO injection in the Gi-DREADD group (n = 4 mice). (M) Proportion of non-bulbous and bulbous microglial endings contacting NE boutons before and after CNO injection in the Gi-DREADD group (n = 4 mice). (N, O) Representative time-lapse images of microglia (GFP) interacting with NE axons (tdTomato) before (N) and after (O) CNO injection in the Gq-DREADD group. Scale bar, 5 μm. (P) Representative two-photon images of microglia (GFP) interacting with NE axons (tdTomato) before (left) and after (right) CNO injection in the Gq-DREADD group. White boxes indicate magnified regions in the right panels; white arrowheads mark bouton contacts. Scale bar, 10 μm; magnified images, 5 μm. (Q, R) Quantification of the number (Q) and duration (R) of microglial process contacts with NE axonal boutons or shafts before and after CNO injection in the Gq-DREADD group (n = 4 mice). (S) Proportion of non-bulbous and bulbous microglial endings contacting NE boutons before and after CNO injection in the Gq-DREADD group (n = 4 mice). Data are presented as mean ± SEM. Statistical significance was determined using a paired t-test in all graphs, *P < 0.05; **P < 0.01; *** P < 0.001; **** P < 0.0001. Each circle indicates an individual mouse.

### Physical contacts between microglial processes and NE boutons

To investigate the structural basis of microglial interactions with NE boutons, we performed super-resolution confocal imaging via the Spinning Disk Super Resolution by Optical Pixel Reassignment system. Microglia were labeled with GFP, and NE axons were identified by immunostaining for dopamine β-hydroxylase (DBH) (Figure 4A). Confocal and 3D reconstructions revealed physical associations between microglial processes and NE axon boutons (Figure 4B). To further examine these interactions at the ultrastructural level, we employed correlative serial block-face scanning electron microscopy (SBF-SEM). NE axons were selectively labeled with tdTomato via AAV-mediated expression (AAV9-TH-Cre combined with AAV9-CAG-FLEX-tdTomato) targeted to the LC, and microglia were labeled with GFP in *Cx3cr1^GFP^*^/+^ mice (Figure 4C). Regions of interest (ROIs) identified by confocal imaging were precisely marked by laser branding under two-photon microscopy^23^. Subsequent 3D reconstructions and SBF-SEM images revealed that microglial process endings directly contact NE boutons without an intervening space (Figure 4D–E). In addition, microglial filopodia, which exhibit fewer contacts with NE axons and are typically classified as non-bulbous endings, were observed making direct contacts with NE boutons as well (Figure S5A-C). Notably, these boutons displayed varicose structures containing small vesicles and formed synaptic contacts with a dendritic spine-like structure (Figure S5D). Together, these findings suggest that both microglial process endings and filopodia physically interact with NE boutons.

**Figure 4.**
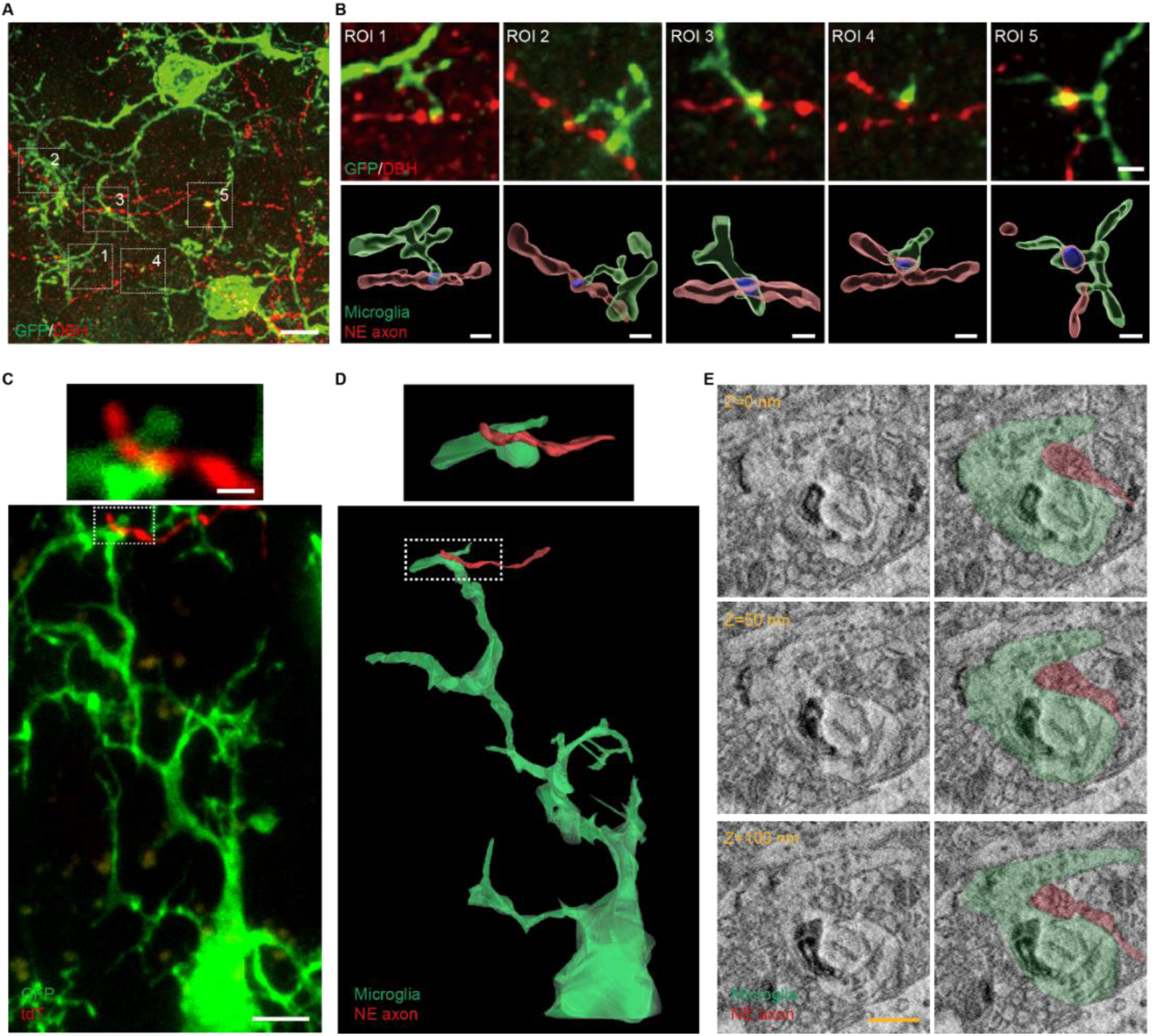
Super-resolution and electron microscopy reconstructions revealed structural features of microglia-NE axon interactions. (A) Representative super-resolution confocal images of GFP-labeled microglia and DBH-labeled NE axons in the cortex. White boxes indicate regions magnified in (B). Scale bar, 5 μm. (B) Magnified images of five ROIs from (A) showing microglia interacting with NE axonal boutons (upper panel) and the corresponding 3D surface renderings generated with Imaris (lower panel). Scale bar, 1 μm. (C) Representative confocal images showing microglia (GFP) with NE axon (tdTomato) interactions in the cortex, followed by electron microscopy reconstruction. White boxes indicate regions magnified in the top panel. Scale bar, 5 μm. Magnified images, Scale bar, 1 μm. (D) 3D serial reconstruction of a microglia (green) with NE axon (red) interaction. White boxes indicate regions magnified in the top panel. (E) Electron micrographs showing a microglial process (green) contacting an NE axon (red) at the ultrastructural level across different Z-planes using SEM. Scale bar, 500 nm.

### Microglial contact with NE axons enhances Ca^2+^ signaling

Next, we tested the functional significance of the interaction between microglia and NE axons in sleep-wake cycle. To this end, we performed two-photon in vivo imaging to monitor Ca^2+^ dynamics in NE axons. We injected AAV2/9-TH-FLP and AAV2/9-hSyn-fDIO-axon-GCaMP6s into the LC of *Cx3cr1^CreER/+^; Ai14* mice (Figure S7A and Video S3). The GCaMP6s reporter was validated by delivering air-puff stimulation to awake mice, which reliably evoked robust Ca^2+^ transients in NE axons within the frontal cortex (Figure S7B–E). As expected, NE axons exhibited significantly higher Ca^2+^ activity during wakefulness than during NREM sleep (Figure S7F–J), consistent with the established role of NE in arousal.

To understand the potential role of microglia-NE axon interaction, we examine Ca^2+^ activity in NE axons relative to microglial proximity in the frontal cortex. Interestingly, NE axons showed a gradual and sustained increase in Ca^2+^ signal intensity when contacted by microglial processes (Figure 5A, B and Video S4). By quantification, 57.5% NE axons show increased Ca^2+^ activity, 39.2% show no changes, and 3.3% show decreased activity after microglial contacts (Figure 5C, D, G). In contrast, after withdrawal of microglial processes, 62.2% of NE axons showed no change in Ca²⁺ signal intensity within the 6 minutes after withdrawal of contacts, whereas 26.3% decreased and 11.5% increased their activity (Figure 5E, F, G). Notably, this modulatory effect of microglia was observed regardless of brain state, as similar increases in Ca^2+^ activity of NE axons upon microglial contact were detected during both wakefulness (Figure 5H, I) and sleep (Figure 5J, K). Together, these findings suggest that microglial contact enhances the Ca^2+^ activity of NE axons in a state-independent manner, indicating a role for microglia–NE axon interactions in modulating noradrenergic tone.

**Figure 5.**
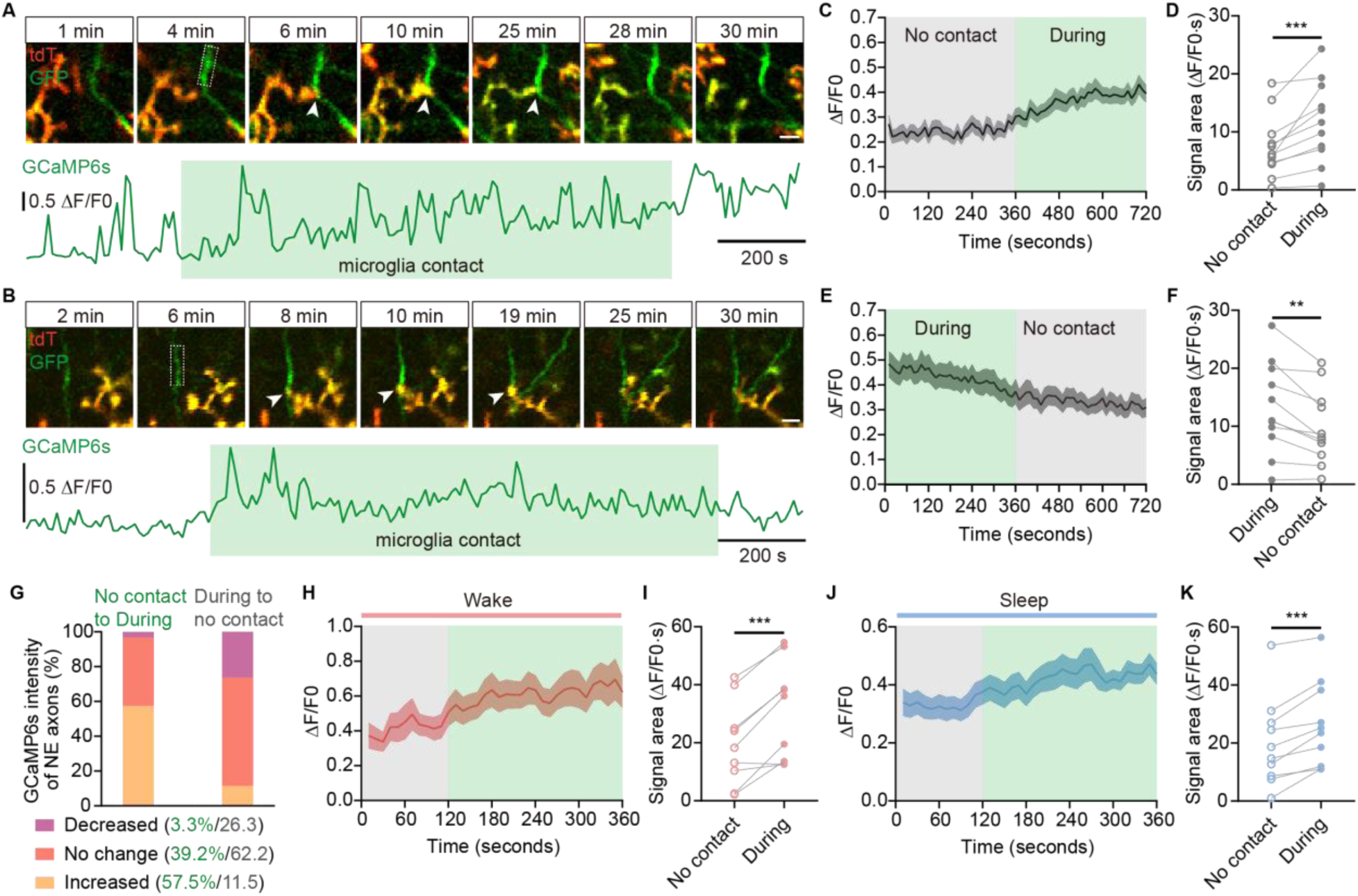
Microglial contact with NE axon enhances Ca^2+^ signaling independent of sleep-wake states. (A, B) Representative time-lapse images showing interactions between microglial processes (tdTomato) and NE axons (GFP), along with corresponding Ca²⁺ activity (ΔF/F0) during 30 min of two-photon imaging. Arrowheads indicate the contact sites. The white box indicates the ROI used for the quantification of calcium signals of NE axons. The green-shaded area indicates the period of microglial contact with the NE axon. Scale bar, 5 μm. (C) Average Ca²⁺ activity (ΔF/F0) across all ROIs of NE axons from no-contact to contact episodes (153 ROIs from 11 mice). (D) Quantification of integrated Ca²⁺ activity (ΔF/F0·s) during no-contact and contact episodes (n = 11 mice). (E) Average Ca²⁺ activity (ΔF/F0) across all ROIs of NE axons from contact to no-contact episodes (142 ROIs from 11 mice). (F) Quantification of integrated Ca²⁺ activity (ΔF/F0·s) during contact and no-contact episodes (n = 11 mice). (G) Proportion of changes in integrated Ca²⁺ activity (ΔF/F0·s) (increase, decrease, no change) induced by microglial contacts and following termination of microglial contacts (n = 11 mice). (H) Average Ca²⁺ activity (ΔF/F0) across all ROIs of NE axons from no-contact to contact in the wake state (48 ROIs from 9 mice). (I) Quantification of integrated Ca²⁺ activity (ΔF/F0·s) during contact and no-contact episodes in the wake state (n = 9 mice). (J) Average Ca²⁺ activity (ΔF/F0) across all ROIs of NE axons from no-contact to contact in the sleep state (127 ROIs from 10 mice). (K) Quantification of integrated Ca²⁺ activity (ΔF/F0·s) during contact and no-contact episodes in the sleep state (n = 10 mice). Data are presented as mean ± SEM. Statistical significance was determined using a paired t-test in all graphs, *P < 0.05; **P < 0.01; *** P < 0.001; **** P < 0.0001. Each circle or point indicates an individual mouse.

### Microglial β2AR mediates microglia-NE axon interaction to sustain wakefulness

Activation of microglial β2AR induces process retraction and reduces microglial surveillance^21,22^. To determine whether NE-β2AR signaling modulates microglial interactions with NE axons, we performed in vivo two-photon imaging combined with EEG–EMG recordings in *Cx3cr1^CreER/+^;Adrβ2^fl/fl^; Ai14* mice. NE axons were labeled with EGFP via viral injection (AAV2/9-TH-FLP mixed with AAV2/9-hSyn-fDIO-EGFP) in the LC region, and microglia were labeled with tdTomato (Figure 6A). We found that β2AR deletion abolished the enhanced microglial surveillance observed during sleep (Figure 6B, C). Furthermore, β2AR deletion eliminated the sleep-induced increase in contacts between microglia and NE boutons (Figure 6D, E), while contact duration remained indistinguishable between sleep and wake states (Figure 6F). Additionally, we chemogenetically manipulated LC neuronal activity and corresponding cortical NE levels in control or microglial β2AR conditional knockout (β2AR cKO; *Cx3cr1Cre^ER/+^; Adrβ2^fl/fl^*) mice via LC-targeted injections of control AAVs (AAV2/9-TH-FLP + AAV2/9-hSyn-fDIO-EGFP), Gq-DREADD (AAV2/9-TH-FLP + AAV2/9-hSyn-fDIO-hM3D(Gq)-EGFP), or Gi-DREADD (AAV2/9-TH-FLP + AAV2/9-hSyn-fDIO-hM4D(Gi)-EGFP) (Figure S7A). Immunostaining analysis revealed that loss of microglial β2AR abolished the complexity changes normally observed in response to fluctuations in cortical NE levels (Figure S7B–D). Consistent with in vivo data, microglial contacts with NE axons did not differ significantly among control, Gq-DREADD, and Gi-DREADD groups in β2AR cKO mice (Figure S7E–H). Thus, these findings suggest that NE–β2AR signaling modulates microglia–NE axon interactions by altering microglial surveillance.

**Figure 6.**
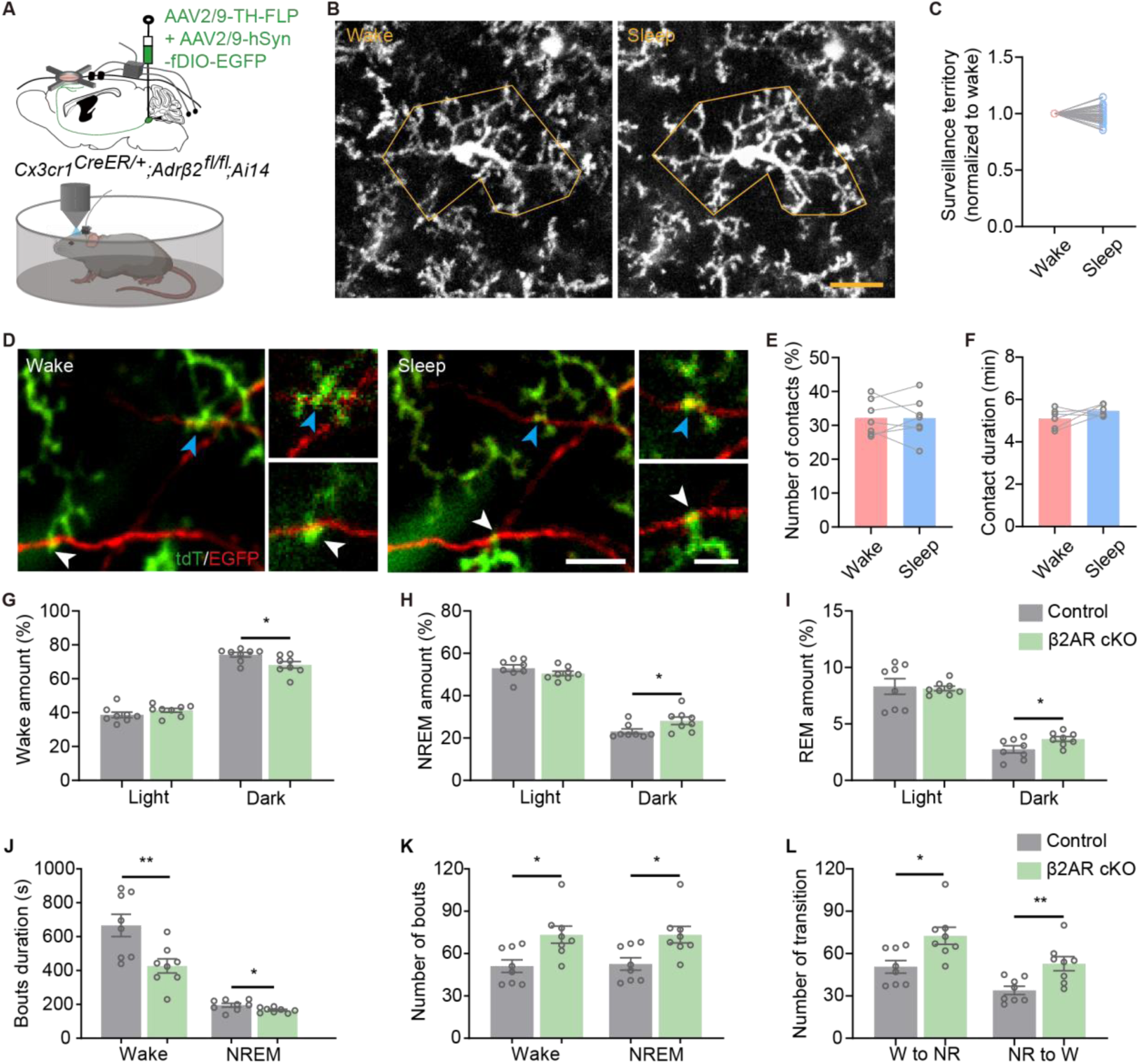
Microglial β2AR mediates microglia-NE axon interaction to sustain wakefulness. (A) Experimental design for examining microglial interactions with NE axonal boutons in β2AR cKO mice during wake and sleep periods using head-fixed two-photon imaging. (B) Representative two-photon images of microglia (max intensity projection) during wake and sleep periods. The yellow outline indicates the surveillance territory of the microglial process over a 10-minute period during wake and sleep states. Scale bars, 10 μm. (C) Quantification of microglial occupied area during 10-minute wake and sleep periods (n=6 mice). (D) Representative two-photon images of microglia (tdTomato) interacting with NE axons (EGFP) during wake and sleep periods. Arrowheads mark microglial process endings in contact with NE axonal bouton; magnified regions are shown in the right panels. Scale bar, 10 μm; magnified images, 5 μm. (E, F) Quantification of the number (E) and duration (F) of microglial process endings in contact with NE axonal boutons during wake and sleep (n = 6 mice). (G–I) Percentage of time spent in wakefulness (G), NREM sleep (H), and REM sleep (I) during the light and dark phases in control and β2AR cKO mice (Control and β2AR cKO group, n=8 mice). (J–L) Quantification of sleep architecture during the dark phase, including bout duration (J), number of bouts (K), and number of transitions between wakefulness and NREM sleep (L) in control and β2AR cKO mice (Control group, n=8 mice; β2AR cKO group, n=8 mice). Data are presented as mean ± SEM. Statistical significance was determined using a paired t-test in C, E, F; an unpaired t-test in G-L; *P < 0.05; **P < 0.01; *** P < 0.001; **** P < 0.0001. Each circle indicates an individual mouse.

To further understand microglia β2AR signaling in regulation of sleep-wake cycle, we examined the sleep pattens in mice deficient of β2AR. We fond that β2AR cKO mice reduced wakefulness and elevated NREM and REM sleep during the dark phase compared with control mice (Figure 6G-I). In addition, β2AR cKO mice show decreased wake and NREM sleep bout duration, accompanied by an increase in the number of wake and NREM sleep bouts, as well as an elevated number of transitions between wake and NREM sleep (Figure 6J-L), indicating impaired stability of nocturnal wakefulness. Together, our results suggest that the microglial β2AR sustains wakefulness by governing microglial engagement with NE boutons.

## Discussion

Microglia have increasingly been recognized for their roles beyond classical immune phagocytosis and inflammation. However, how microglia contribute to the regulation of sleep–wake states remains incompletely understood. In this study, we demonstrate that microglial surveillance is enhanced during sleep, accompanied by increased physical interactions with NE axon boutons. Notably, these interactions are associated with elevated Ca^2+^ activity in NE axons, suggesting a potential modulatory role of microglia in NE signaling. Furthermore, we identified that reduced NE levels enhance microglial surveillance via β2AR, priming microglia for axonal interactions (Figure S8). These findings reveal a previously uncharacterized microglia-neuromodulator interaction in maintaining the sleep-wake cycle.

### Microglial process dynamics and their potential roles differ across wakefulness, NREM sleep, and REM sleep

Investigating microglial process dynamics during natural sleep–wake states remains challenging due to technical limitations. In vivo two-photon imaging studies have produced divergent findings: head-fixed mice showed reduced microglial complexity during NREM sleep compared to wakefulness^16^, whereas miniaturized two-photon imaging in freely moving mice revealed increased complexity during both NREM and REM sleep^37^. These discrepancies likely reflect differences in experimental conditions and analytical approaches. Miniature imaging in freely moving mice provides a more natural representation of sleep–wake states, with fewer microarousals and more stable NREM sleep, whereas head restraint, even after habituation^38^, may induce stress responses that influence microglial morphology and dynamics^39^.

In our study, we started with continuous two-photon imaging in head-fixed mice for 3 hours following a ten-day habituation period. Representative 10-minute epochs of sleep and wakefulness during the daytime were selected based on EEG-EMG scoring. Although brief state transitions occurred, each epoch consisted of over 80% of the target state, ensuring overall consistency. Within these windows, we observed significantly enhanced microglial surveillance during NREM sleep. To confirm these findings, we repeated key experiments using miniaturized two-photon imaging in freely moving mice and applied the same analytical pipeline, again observing increased microglial surveillance during sleep. Due to the short duration of daytime REM episodes (averaging less than 2 minutes during the daytime)^40^, microglial dynamics during REM were not directly assessed in our study.

REM sleep is initiated by REM-on neurons in the brainstem and maintained through a balance of excitatory cholinergic drive and suppressed monoaminergic tone^32^. Given that NREM and REM sleep are regulated by distinct brain regions, microglial process dynamics may likewise exhibit region– and state-specific characteristics. As critical regulators of synaptic structure and function^9,13,15,41^, microglia appear to exert distinct, vigilance-state-dependent effects across sleep stages. Supporting this notion, a study has shown that microglia-induced increases in spine activity are dependent on vigilance state, occurring predominantly during NREM sleep^16^. In contrast, disturbances in REM sleep are associated with increased microglial activation^42,43^, highlighting their potential role in REM regulation and modulation of synaptic plasticity during this sleep stage. Together, these findings suggest that microglia may contribute to sleep stage-dependent synaptic regulation, although further mechanistic studies are needed.

### Microglia promote Ca^2+^ activity of NE axons

Microglia are highly dynamic sensors of neuronal activity and exert diverse modulatory effects on neural circuits. They dampen neuronal activity by converting ATP to adenosine^44^ and extend processes toward active neurons, where their contacts further suppress spontaneous and evoked activity^36^. Microglial contact also shapes structural plasticity, inducing filopodia formation in the developing somatosensory cortex^41^ and supporting spine remodeling essential for learning and memory in adult mice^45^. In addition, microglia have been shown to selectively enhance the activity of synapses and neurons they contact under physiological contexts^15,16^.

Previous studies have shown that microglia can engulf NE fibers both in the trigeminal spinal subnucleus caudalis following infraorbital nerve injury ^46^ and in the olfactory bulb during early AD^47^. These studies highlight an interaction between microglia and NE axons under pathological contexts. The question is how microglia regulate neuromodulatory transmission, such as noradrenergic signaling? Using two-photon in vivo imaging and correlative EM, our current study demonstrates that cortical microglia have direct contact with NE axon boutons and the interactions increase local Ca^2+^ activity at NE axons. Interestingly, our results showed that microglial contact enhanced Ca^2+^ activity independent of sleep-wake states. Considering LC neuronal activity regulates vigilance states^34,35^, our results suggest that the local Ca^2+^ elevation at LC axons is induced by physical contact with microglia, likely separated from somatic spiking activity.

The mechanism underlying the increase in NE axon activity following microglial interactions remains unclear. Microglial processes can invade the synaptic cleft to shield GABAergic inputs or displace inhibitory synapses from neurons, thereby reducing inhibitory GABAergic activity^23,48^. Our EM data reveal direct physical contacts between microglial processes and NE axons, which may interfere with inhibitory inputs or α2 receptors on NE axons, potentially enhancing axon Ca^2+^ activity. In addition, cell adhesion molecules (CAMs), which are highly expressed on neurons and glial cells, play critical roles in synaptic plasticity, axon–myelin interactions and cell–cell communication^49–51^. Activation of CAMs, including neural cell adhesion molecule 2, can elevate intracellular Ca²⁺ levels by triggering diverse signaling cascades or through L-type voltage-dependent calcium channels in developing cortical neurons^52,53^. Such CAM-mediated mechanisms by direct microglial contact may potentially contribute to the increased Ca²⁺ activity observed in NE terminals following microglial interactions.

### Microglial β2AR signaling regulates the sleep-wake cycle

Extracellular NE serves as a crucial modulator of microglial surveillance and motility^21,22^. Microglia express high levels of β2AR, and their activation—either by endogenous NE or pharmacological agonists—induces process retraction and reduces microglial responses to local injury^21,22,54,55^. Conversely, blockade of β2AR increases microglial process surveillance in awake mice^21,22^. In addition, microglial β2AR deficiency diminishes the anesthesia-induced increase in microglial surveillance and reduces bulbous ending contacts with neurons, leading to attenuated neuronal hyperactivity post-anesthesia^22,23^. Together, these findings highlight that microglial β2AR is a key regulator of microglial process dynamics responses to NE and contributes to microglia–neuron interactions.

In our study, we found that cortical NE levels fluctuate dynamically across the sleep–wake cycle. In addition, NE levels regulate both microglial surveillance and microglia–NE axon interactions through β2AR, thereby enhancing local Ca²⁺ signaling of NE axons. Synaptic strength is closely linked to presynaptic Ca^2+^ dynamics^56,57^, and microglial process–induced Ca^2+^ elevations at NE axons likely facilitate NE release. Considering the critical role of cortical NE in maintaining wakefulness, we propose that this sleep-associated NE release, triggered by microglial contact, serves to prime the brain for subsequent wakefulness by promoting the maintenance of arousal. In line with this, β2AR cKO mice exhibit disrupted microglia–NE axon interactions accompanied by reduced wake stability during the dark phase.

Previous work reported that global β2AR knockout mice exhibit reduced wake time and increased REM sleep. It also abolished differences in microglial dynamics between REM and NREM sleep but not between wake and NREM states, while ablation of LC-NE projections eliminated state dependence altogether^37^. These findings suggest that NE signaling through β2AR contributes partially, but not exclusively, to sleep-state regulation of microglial activity. Here, we found that microglial-specific β2AR deficiency diminishes their ability to adjust morphological complexity and microglia-NE axon interactions across the sleep-wake cycle or varying NE levels. Moreover, microglial β2AR deficiency led to altered sleep architecture, characterized by reduced wakefulness and increased NREM and REM sleep during the dark phase. The differences between global and conditional β2AR models may reflect the broader impact of β2AR loss on non-microglial cell types that indirectly influence microglial dynamics during the sleep-wake cycle. Nevertheless, our current study highlights the critical role of NE-β2AR signaling in microglia–axon interactions and its contribution to promoting the maintenance of wakefulness in the sleep-wake cycle.

## Methods

### Animals

Animal care and procedures were approved by the University of Texas Health Science Center Houston Institutional Animal Care and Use Committee. Mice were housed at 21–22 °C on a 12 h light/12 h dark cycle with standard pellet chow and water ad libitum otherwise noted for PLX3397 and control diet treatment. Both female and male mice aged 6–10 weeks were used for all experiments. *C57BL6/J (JAX:000664), Cx3cr1^GFP/GFP^ (B6.129P2(Cg)-Cx3cr1^tm1Litt^/J, 005582), R26^hM4Di/hM4Di^* (*B6.129-Gt(ROSA*)*26Sor^tm1(CAG-CHRM4*,-mCitrine)Ute^*/*J*, *026219*), *R26^hM3Dq/mCitrine^* (*B6N;129-Tg(CAG-CHRM3*,-mCitrine)1Ute/J, 026220), Cx3cr1^CreER/CreER^ (B6.129P2(Cg)-Cx3cr1tm2.1(cre/ERT2)Litt/WganJ, 021160), Ai14 (B6.Cg-Gt(ROSA)26Sortm14(CAG-tdTomato)Hze/J, 007914)* mouse lines were acquired from the Jackson Laboratory and subsequently bred in the animal facility at UTH. *Cx3cr1^CreER/CreER^* mice were crossed with *Ai14* reporter mice to generate offspring in which microglia were fluorescently labeled with tdTomato. *Adrβ2^fl/fl^*mice (generously provided by G. Karsenty at Columbia University) were subsequently crossed with *Cx3cr1^CreER/CreER^* and *Ai14* mice to produce animals in which the β2AR was selectively knocked out in microglia.

### Surgery

For all survival surgeries, adult mice were given ibuprofen (0.2 mg/ml) in their drinking water for 3 days prior to the procedure. Anesthesia was induced with 4%–5% isoflurane in an induction chamber and maintained at 1.5%–2% isoflurane throughout the surgery. Mice were placed in a stereotaxic frame, the head was secured, and the scalp was shaved and incised to expose the skull.

#### Cranial window implantation

For cranial window implantation, a circular region (3.5 mm in diameter) over the unilateral frontal cortex was drilled using a high-speed microdrill. The skull within this area was carefully removed, and a sterilized glass coverslip (3.5 mm diameter; Warner Instruments) was placed over the craniotomy and secured using light-curing dental cement (Tetric EvoFlow or equivalent). Once the cement was cured, the remaining exposed skull (excluding the window region) was coated with iBond Total Etch primer (Heraeus) and cured with an LED light. A sterilized stainless steel headplate was then affixed to the skull using light-curing dental cement applied over the iBond primer layer.

#### EEG–EMG electrode placement

For experiments requiring simultaneous EEG–EMG recordings, electrode implantation was performed immediately following cranial window placement. After securing the glass coverslip and coating the remaining skull with iBond Total Etch primer, two stainless steel screws were inserted into the frontal and lateral parietal regions within the same hemisphere for EEG signal acquisition. Two additional screws were placed in the lateral parietal region for grounding. Two silver EEG wires and ground wires were tightly wrapped around the corresponding screws. For EMG recording, two insulated silver wire electrodes were inserted into the neck muscles. The EEG–EMG electrode assembly was secured over the interparietal region and affixed using light-curing dental cement. The headplate was also attached using light-curing dental cement.

#### Virus injection through cranial window

For experiments requiring viral injections within the cranial window area, AAVs were injected immediately following the removal of the skull over the target region. Injections were performed into the center of the cranial window area of the frontal cortex (AP: +1.7 mm; ML: +1.5 mm; DV: −0.5 mm) using a glass micropipette connected to a microinjection pump (R-480, RWD Life Science, Shenzhen, China). The injection rate was maintained at 35 nl/min. Following the injection, the pipette was left in place for an additional 5 minutes to diffuse widely, and then slowly retracted.

To monitor norepinephrine (NE) dynamics, AAV2/9-hSyn-NE2h (rAAV-hSyn-NE2h-WPRE-hGH polyA, AAV2/9; BrainVTA, PT-5262) was used. For ATP signal monitoring, AAV2/9-hSyn-ATP1.0 (rAAV-hSyn-ATP1.0-WPRE-hGH pA, AAV2/9; BrainVTA, PT-1350) was employed. In both cases, approximately 400 nl of virus was injected into the frontal cortex.

#### Virus injection in the locus coeruleus region

For experiments requiring viral injections in the locus coeruleus (LC) region, following scalp incision and skull exposure, two small craniotomies were made above the target sites. AAVs were then bilaterally injected into the LC at the following coordinates: AP = −6.00 mm, ML = ±0.95 mm, DV = −3.12 mm.

To label NE axons in the cortex, 150 nL of a 1:3 volume mixture of AAV9-TH-Cre (AAV.rTH.PI.Cre.SV40; Addgene, 107788-AAV9) and AAV-CAG-FLEX-tdTomato (pAAV-FLEX-tdTomato; Addgene, 28306-AAV9) was slowly injected into the bilateral LC of *Cx3cr1^GFP/+^*mice at a rate of 20 nL/min.

To monitor Ca^2+^ activity in NE axons of the frontal cortex, 150 nL of a 1:3 volume mixture of AAV2/9-TH-FLP (rAAV-TH-FLP-WPRE-hGH pA, AAV2/9; BrainVTA, PT-2973) and AAV2/9-hSyn-fDIO-axon-GCaMP6s (rAAV-hSyn-fDIO-axon-GCaMp6s-WPRE-hGH polyA, AAV2/9; BrainVTA, PT-9272) was injected (20 nL/min) into the bilateral LC regions of *Cx3cr1^CreER/+^; Ai14* mice.

For chemogenetic manipulation, AAV9-TH-Cre (AAV.rTH.PI.Cre.SV40; Addgene, 107788-AAV9) was injected into the bilateral LC regions of *Cx3cr1^GFP/+^*, *Cx3cr1^GFP/+^:R26^hM3Dq/+^*, and *Cx3cr1^GFP/+^:R26^hM4Di/+^* mice.

For chemogenetic manipulation in the β2AR cKO mice, the bilateral LC region was injected with control AAVs (AAV2/9-TH-FLP + AAV2/9-hSyn-fDIO-EGFP), Gq-DREADD (AAV2/9-TH-FLP + AAV2/9-hSyn-fDIO-hM3D(Gq)-EGFP), or Gi-DREADD (AAV2/9-TH-FLP + AAV2/9-hSyn-fDIO-hM4D(Gi)-EGFP).

To investigate microglia-NE axon bouton interactions in β2AR cKO mice, 150 nL of a 1:3 volume mixture of AAV2/9-TH-FLP (rAAV-TH-FLP-WPRE-hGH pA, AAV2/9; BrainVTA, PT-2973) and AAV2/9-hSyn-fDIO-EGFP (rAAV-hSyn-fDIO-EGFP-WPRE-hGH polyA, AAV2/9; BrainVTA, PT-3622) was injected (20 nL/min) into the bilateral LC of *Cx3cr1^CreER^; Adrβ2^fl/fl^; Ai14* mice.

All viral preparations were standardized to a final working titer of 1–5 × 10¹² vg/mL.

#### Optical fiber and EEG–EMG electrode implantation

Following virus injection (AAV2/9-hSyn-NE2h or AAV2/9-hSyn-ATP1.0) into the frontal cortex (AP: +1.7 mm; ML: +1.5 mm; DV: −0.5 mm), an optical fiber was implanted 0.2 mm above the injection site. The fiber was secured to the skull using dental cement. Subsequently, stainless steel screws were inserted into the frontal and lateral parietal regions, and EEG and ground wires were tightly wrapped around the respective screws as previously described. Finally, the exposed skull was completely covered, and EEG-EMG electrode was secured using light-curing dental cement.

### Drug administration

To induce CreER-mediated recombination, tamoxifen (MilliporeSigma, catalog no. T5648) was administered via intraperitoneal (i.p.) injection 6 weeks before imaging (150 mg kg⁻¹, dissolved at 20 mg ml⁻¹ in corn oil; five injections in total, spaced 48 h apart) in *Cx3cr1^CreER/+^; Ai14* and *Cx3cr1^CreER/+^; Adrβ2^fl/fl^; Ai14*mice. To pharmacologically deplete microglia, mice were fed chow containing PLX3397 (a CSF1 receptor inhibitor) for 3 weeks (600 mg kg⁻¹; Chemgood, catalog no. C-1271). Animals were randomly assigned to different treatment groups.

For chemogenetic manipulation, Clozapine N-oxide (Cayman Chemical, 12059) was administered via i.p. injection at a dose of 1.5 mg/kg to mice in both the Gi-DREADD and Gq-DREADD groups, as well as to control mice.

### EEG-EMG recording and data analysis

For 24-hour EEG–EMG recordings in both the microglia-ablated and control groups, as well as the β2AR cKO and control groups, animals were allowed to recover for at least one week following EEG–EMG electrode implantation. They were then placed in the recording cage and habituated to the recording cables and environment for 1 day.

For recordings paired with two-photon imaging (either head-fixed or freely moving), EEG–EMG recording was initiated prior to the imaging session. The start time of imaging was marked in the EEG–EMG data file for alignment.

Cortical EEG and EMG signals were acquired at a sampling rate of 250 Hz using the Medusa system (Bio-Signal Technologies, Nanjing, China). Data were uploaded to the Lunion online scoring platform (https://stage.luniondata.com), where polygraphic recordings were automatically segmented into wakefulness, NREM sleep, or REM sleep in 4-s or 10-s epochs according to standard criteria. For each epoch, delta (0.5–4 Hz), theta (4–10 Hz), and sigma (10–15 Hz) band powers were calculated from the EEG, and an EMG envelope was obtained by rectification followed by 10-Hz low-pass filtering. Wakefulness was defined as high EMG activity with low delta power; NREM sleep as high delta power with low-to-moderate EMG activity; and REM sleep as muscle atonia (low EMG) with an elevated theta/delta power ratio. All automatically assigned sleep–wake stages were visually inspected and manually corrected when necessary.

### In vivo two-photon imaging in head-fixed mice

Mice were allowed to recover for two weeks following cranial window implantation. They were then trained to move freely on an air-lifted platform (Mobile HomeCage, Neurotar Ltd., Helsinki, Finland) while head-fixed under a two-photon microscope. To reduce the stress induced by head restraint in mice, we habituated them to the head-fixed condition for 10 days prior to the start of imaging (*PMID: 32699235*). During the first 3 days, mice were habituated to head fixation, beginning with 30-minute sessions and gradually increasing to 1 hour per day. Over the subsequent 7 days, mice were head-fixed under the two-photon microscope for 1 to 3 hours per day at randomly assigned durations. To promote stable and sustained sleep episodes, both habituation and imaging were performed during the light phase, primarily between 2 to 5 hours after lights-on.

Mice were imaged using a two-photon microscope system (Scientifica Ltd., Uckfield, UK) equipped with a tunable Ti: Sapphire Mai Tai DeepSee laser (Spectra-Physics, Santa Clara, CA, USA). The laser wavelength was set to 920 nm for imaging GFP and EGFP signals, or to 940 nm for simultaneous imaging of tdTomato and GFP/EGFP signals. A 16× water-immersion objective (Nikon) was used, with digital zoom (2x to 5x) applied as needed depending on experimental requirements. The microscope was equipped with 525/50 nm and 620/60 nm emission filters for detecting GFP (green channel) and tdTomato (red channel), respectively. Laser power at the sample was maintained between 30 and 40 mW. Imaging of the cortex was performed at depths ranging from 70 to 200 μm below the pial surface.

To investigate microglial process dynamics and their interactions with NE axons across the sleep–wake cycle and during chemogenetic manipulation, a combination of z-stack and time-series imaging was performed. For z-stack imaging, a total depth of 20 μm was scanned with 2 μm step intervals, with each complete z-stack loop taking 1 minute. Time-series imaging was used specifically to monitor NE and axon GCaMP6s signals. Each imaging session lasted up to 3 hours and was performed concurrently with EEG–EMG recording. Imaging during chemogenetic manipulation lasted approximately 35 minutes for each session.

### In vivo two-photon imaging in freely moving mice

Following cranial window implantation, mice were allowed to recover for two weeks. Prior to imaging, mice were head-fixed on a treadmill to locate the imaging region and attach the basement for fixing the probe. Mice were then housed in the recording cage for one day to allow environmental acclimation before the imaging session began.

On the day of imaging, the miniature microscope probe was inserted into the basement, and the EEG-EMG recording electrodes were connected to the recording system.

For two-photon microscopy of tail suspension, a miniature two-photon imaging system (SUPERNOVA-100, headpiece: FHIRM-U; Transcend Vivoscope Biotech Co., Ltd., China) was employed. Imaging data were acquired using imaging software (SUPERGIN, Transcend Vivoscope Biotech Co., Ltd., China) at a frame rate of 5 Hz (1024 × 880 pixels) with a 920 nm femtosecond fiber laser. The average power delivered to the brain was less than 60 mW. Similar to conventional head-fixed two-photon imaging, we captured xyzt imaging to assess microglial process dynamics during the sleep–wake cycle. Imaging sessions were conducted during the daytime.

### Two-photon imaging data analysis

Z-stack and time-series images acquired on a single channel (e.g., microglia-GFP) were corrected for focal plane displacement using ImageJ (NIH, https://imagej.nih.gov/ij/) with the TurboReg and StackReg plugins. For dual-channel images (e.g., microglia-GFP and NE axon-tdTomato), the HyperStackReg plugin was used for alignment and correction.

To assess microglial surveillance territory, Z-projections of average intensity (using every 6-11 slices) were generated to capture the full morphology of microglia. For quantifying the microglia-occupation, maximum intensity projections of time-series images were used for both wake and sleep phases.

To assess microglia–NE axon interactions, Z-projections of average intensity (using every 3-4 slices) were generated, and we manually quantified the number of contacts between individual microglia and NE axons during 10-minute wake and sleep periods, as well as before and after CNO injection. For the latter, 10-minute epochs were randomly selected from 30-minute time-lapse imaging sessions conducted pre– and post-CNO in the chemogenetic manipulation groups.

For microglial contacts with NE axon boutons or shafts, regions of interest (ROIs) were manually drawn in the boutons and axon shafts with consistent signal across the imaging session (10-minute epochs during wake, sleep, and pre– and post-CNO injection). Microglial process contacts were then quantified based on the number of microglial endings contacting these ROIs.

To assess the coverage of microglia (GFP signal) within NE axons (tdTomato signal), NE axons were manually segmented as ROIs using the Segmented Line tool in ImageJ. GFP intensity within these NE ROIs was measured, and the proportion of segments with GFP intensity greater than 0.25 was compared between wake and sleep phases.

To assess Ca^2+^ activity in NE axons, average intensity projections were generated every 10 frames. NE axons were manually segmented as regions of interest (ROIs) using the Segmented Line tool in ImageJ. To minimize fluorescence bleed-through from microglia (tdTomato) into the GCaMP6s signal, ROIs were selected adjacent to, but not directly overlapping with, sites of microglial process contact. For each ROI, mean fluorescence intensity was calculated and expressed as ΔF/F0 = (F – F0) / F0, where F0 was attained from the background fluorescence of each frame. GCaMP6s fluorescence intensity within these NE segments was quantified before and after air puff stimulation, during both sleep and wake states. Additionally, Ca^2+^ activity was compared between NE segments with and without microglial contacts. To classify changes in Ca^2+^ activity before and after microglial contact, the following criteria were applied: increased activity was defined as a post-contact signal exceeding 1.5 times the baseline, while decreased activity was defined as a signal falling below 50% of the baseline. The calcium signal area (ΔF/F0·s) represents the sum (integral) of all ΔF/F0 values exceeding 0.2.

To measure NE dynamics across the sleep-wake cycle, the mean fluorescence intensity of EGFP from one imaging field was calculated and expressed as ΔF/F0 = (F – F0) / F0, where F0 was defined as the 25th percentile (lower quartile) of fluorescence values across all frames.

### Fiber photometry recording and data analysis

Three weeks after surgery for optical fiber and EEG-EMG electrode implantation, mice were placed in the recording cage for one day of environmental habituation. NE signals from the frontal cortex were recorded using a fiber photometry system (Inper Ltd., China). The EGFP-based NE biosensor was excited using a 470 nm LED, with laser output adjusted to 40 μW at the fiber tip. Fluorescence signals were collected at a sampling rate of 20 Hz with an exposure time of 10 ms. Each recording session lasted 2 hours to capture sufficient episodes of wakefulness, NREM sleep, and REM sleep.

Data were analyzed using Inper Plot software (Inper Ltd., China). Fluorescence signals excited by 470 nm light were used for analysis. The raw data were smoothed using a moving average filter with a window size of 5. Baseline correction was performed using a second-order polynomial fit. The fluorescence change was calculated as ΔF/F0 = (F – F0) / F0, where F0 represents the average fluorescence signal over the entire recording period.

### Immunohistochemistry and data quantification

Following completion of behavioral experiments or imaging, mice were anesthetized with isoflurane and transcardially perfused with approximately 25 mL of phosphate-buffered saline (PBS), followed by ∼25 mL of ice-cold 10% formalin. Brains were carefully extracted, post-fixed in 10% formalin at 4°C for 12 hours, and then cryoprotected in 30% sucrose at 4°C overnight. Coronal brain sections (35 μm thick) were cut using a freezing microtome (CM1950, Leica, Wetzlar, Germany). For immunohistochemical staining, sections were washed in PBS (3 × 10 min), then incubated in blocking solution containing 10% Triton X-100 and 5% normal bovine serum in PBS for 60 minutes at room temperature. Subsequently, sections were incubated overnight at 4°C with primary antibodies diluted in antibody buffer containing 0.3% Triton X-100 and 1% normal bovine serum: anti-IBA1 (1:500, Abcam, ab5076), anti-IBA1 (1:500, Wako, 011-27991), anti-DBH (1:500, Abcam, ab209487), anti c-fos (1:500, Abcam, ab190289), anti TH (1:500, Sigma, T1299) and anti-P2Y12 (1:500, AnaSpec, AS-55043A). The following day, sections were washed in PBS (3 × 10 min) and incubated with appropriate secondary antibodies (anti-goat Alexa 488, anti-rabbit Alexa 647, anti-rabbit Alexa 594, anti-goat Alexa 647; anti-mouse Alexa 594, all at 1:500) for 2 hours at room temperature in the dark. After a final PBS wash, sections were mounted on microscope slides and coverslipped with DAPI Fluoromount-G (Southern Biotech, 0100-20).

Imaging was performed using either a Nikon AXR Confocal Laser Microscope with a 60x oil-immersion objective for confocal imaging or a Nikon CSU-W1 SoRa super-resolution spinning disk system for super-resolution imaging.

For microglial morphological analysis based on IBA1 immunostaining, Z-projected images were first segmented using the Trainable Weka Segmentation plugin in ImageJ. After segmentation, 4–8 microglia per mouse were randomly selected for Sholl analysis to assess the complexity of microglial processes.

To assess the interaction between microglia and NE axons, we quantified the colocalized area and counts of IBA1⁺ microglia and DBH⁺ NE fibers using confocal imaging (2,048 × 2,048 pixels; 0.066 μm/pixel; Z-step: 0.6 μm). An average intensity projection was generated by averaging five adjacent optical sections (total thickness: 2.4 μm). Thresholding was applied separately to the IBA1⁺ and DBH⁺ channels to create binary masks. The total area of DBH⁺ signal, as well as the area and number of regions colocalized with IBA1⁺ signal, were quantified to assess microglial interactions with NE axons.

Super-resolution images (2,048 × 2,048 pixels; 0.038 μm/pixel; Z-step: 0.2 μm) obtained from the Nikon CSU-W1 SoRA Spinning Disk confocal microscope were visualized using Imaris to illustrate the three-dimensional interactions between microglial processes and NE boutons.

### Serial block-face scanning electron microscopy (SBF-SEM) and 3D reconstruction

To precisely locate the regions of interest (ROIs) where microglia interact with NE boutons, we employed near-infrared branding (NIRB) as our previously study described^23^. NE axons were labeled by injecting viruses (AAV9-TH-Cre + AAV-CAG-FLEX-tdTomato) into the LC of *Cx3cr1^GFP/+^* mice. Following transcardial perfusion with 4% paraformaldehyde (PFA), brains were coronally sectioned at a thickness of 100 μm using a vibratome, and sections were mounted on glass slides and imaged using confocal microscopy to identify and acquire z-stack images of the regions of interest (ROIs). Next, the slices were transferred to the two-photon microscope, and the previously identified ROIs were relocated. Linear NIRB marks were generated surrounding the target areas using a two-photon laser to facilitate later localization in electron microscopy. After NIRB, the sections were post-fixed in a solution containing 2% glutaraldehyde and 2% paraformaldehyde in 0.1 M cacodylate buffer supplemented with 2 mM Ca2+ chloride at 4 °C for at least 24 hours. The samples were then processed, embedded, and trimmed for SBF-SEM as previously described (*PMID: 38177340*). Serial imaging and sectioning were performed using a VolumeScope 2 scanning electron microscope (Thermo Fisher Scientific) at a resolution of 5 nm × 5 nm × 50 nm (x, y, z).

Electron microscopy image stacks were stitched and smoothed using ImageJ. For three-dimensional (3D) reconstruction, the images were imported into the Reconstruct software package. Based on alignment with confocal image data, the somata of target microglial cells were identified, and both the somata and processes were manually traced throughout the serial sections. Special attention was paid to sites where microglial processes appeared to be in contact with NE axons. NE boutons were identified in each section based on their morphological and ultrastructural features, including small, synaptic vesicles and frequent mitochondrial inclusions (*PMID: 21048893*). All segmented structures were subsequently rendered and visualized in 3D.

### Statistical Analysis

All analyses were conducted using GraphPad Prism (version 10.0, GraphPad Software). Data are presented as mean ± standard error of the mean (SEM) except where otherwise indicated. Normality of data distribution was assessed using the Shapiro–Wilk test. When comparing two time points from the same animal, paired two-tailed t-tests were employed. For comparisons between two independent groups, unpaired two-tailed Student’s t-tests were used for normally distributed data, while the Mann–Whitney U test was applied for non-parametric data. For comparisons involving multiple groups, one-way ANOVA followed by Tukey’s or Dunn’s post hoc tests was performed. Sample sizes and the specific statistical tests applied are provided in the figure legends for each experiment. Statistical significance was set at p < 0.05.

## Acknowledgments

This work is supported by the National Institutes of Health (R35NS132326 and R01NS088627 to L.-J.W.). We thank Mayo Clinic Microscopy and Cell Analysis Core facility for experimental and technical support. We thank the members of the Wu laboratory at UTHealth Houstonfor their insightful discussions.

## Author contributions

Conceptualization and methodology: L.-J.W. and Y.L.; writing – original draft: Y.L. and L.-J.W.; writing – review & editing: L.-J.W. and X. H.; investigation: Y.L. performed most of the experiments and data analysis; W.S. assisted with part of the data acquisition and analysis; E.D. assisted with EEG-EMG and immunostaining data acquisition and analysis; S.Z. and K.H. assisted with electron microscopy data acquisition and analysis; Q.H. assisted with miniature two-photon imaging experiment; Q.H., L.H., and F.Q. assisted with data analysis, resources: L.-J.W.; funding acquisition: L.-J.W.; supervision: L.-J.W.; project administration: L.-J.W. All authors reviewed and approved the final version of the manuscript.

## Competing interests

The authors declare that they have no competing interests.

## Data and materials availability

All data needed to evaluate the conclusions in the paper are present in the paper and/or the Supplementary Materials.

## Supplementary Files

**Figure S1.**
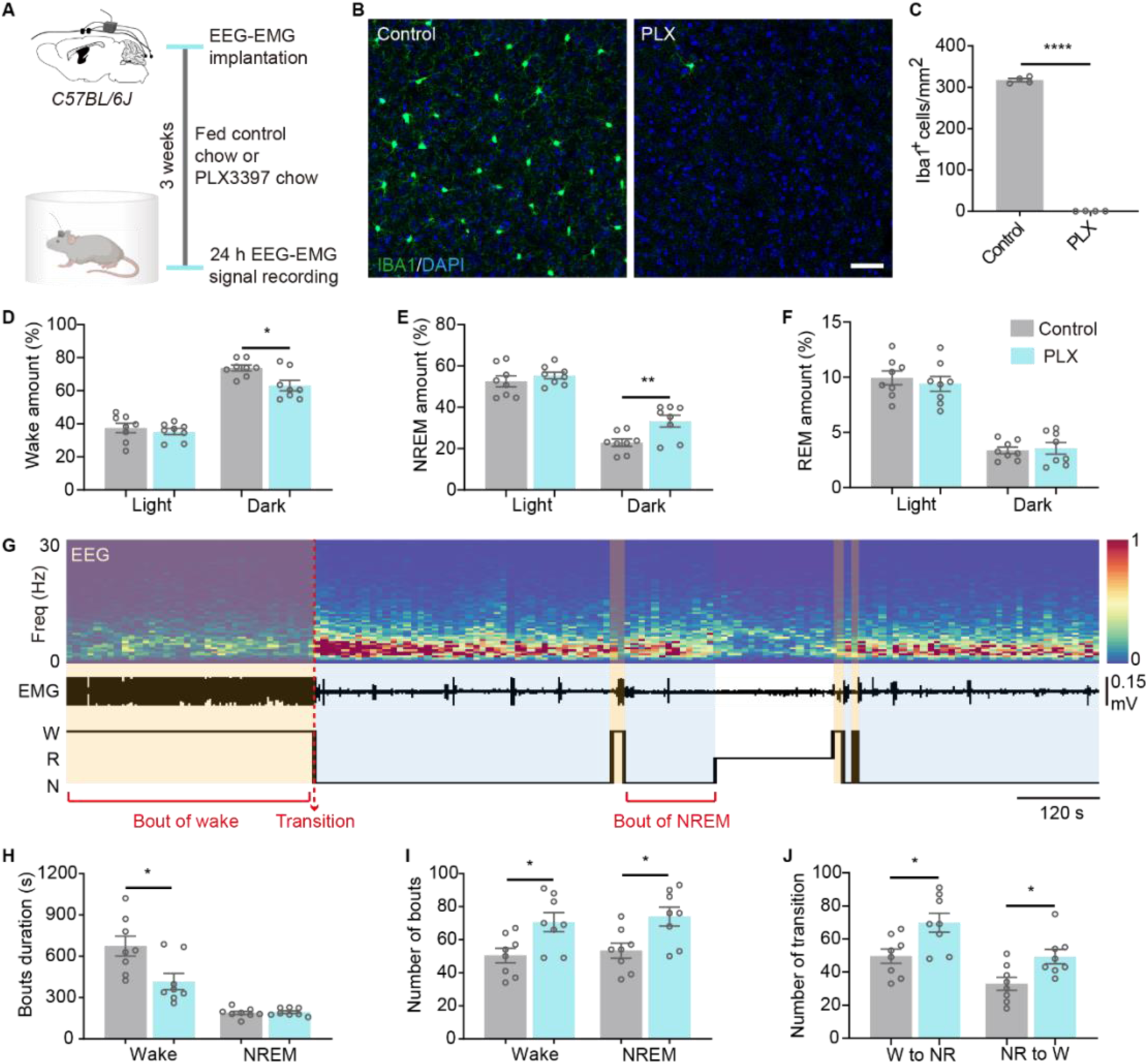
Microglia ablation decreases stable wakefulness during the night. (A) Schematic of the experimental procedure for microglial ablation and EEG–EMG recording. (B, C) Representative immunohistochemical images (B) and quantification (C) of cortical IBA1 and DAPI staining in control and PLX-treated groups (B, Scale bars, 50 μm; C, n=4 mice per group). (D–F) Percentage of time spent in wakefulness (D), NREM sleep (E), and REM sleep (F) during the light and dark phases in control and PLX-treated mice (n=8 mice per group). (G) Representative EEG power spectrum, EMG trace, and hypnogram from a single mouse. Yellow and grey shading indicate bouts of wakefulness and NREM sleep, respectively. The red dashed line marks the transition circle between wake and NREM sleep, or vice versa. (H–J) Quantification of sleep architecture during the dark phase, including bout duration (H), number of bouts (I), and number of transitions between wakefulness and NREM sleep (J) in control and PLX-treated mice (n=8 mice per group). Data are presented as mean ± SEM. Statistical significance was determined using an unpaired t-test in all graphs, *P < 0.05; **P < 0.01; *** P < 0.001; **** P < 0.0001. Each circle indicates an individual mouse.

**Figure S2.**
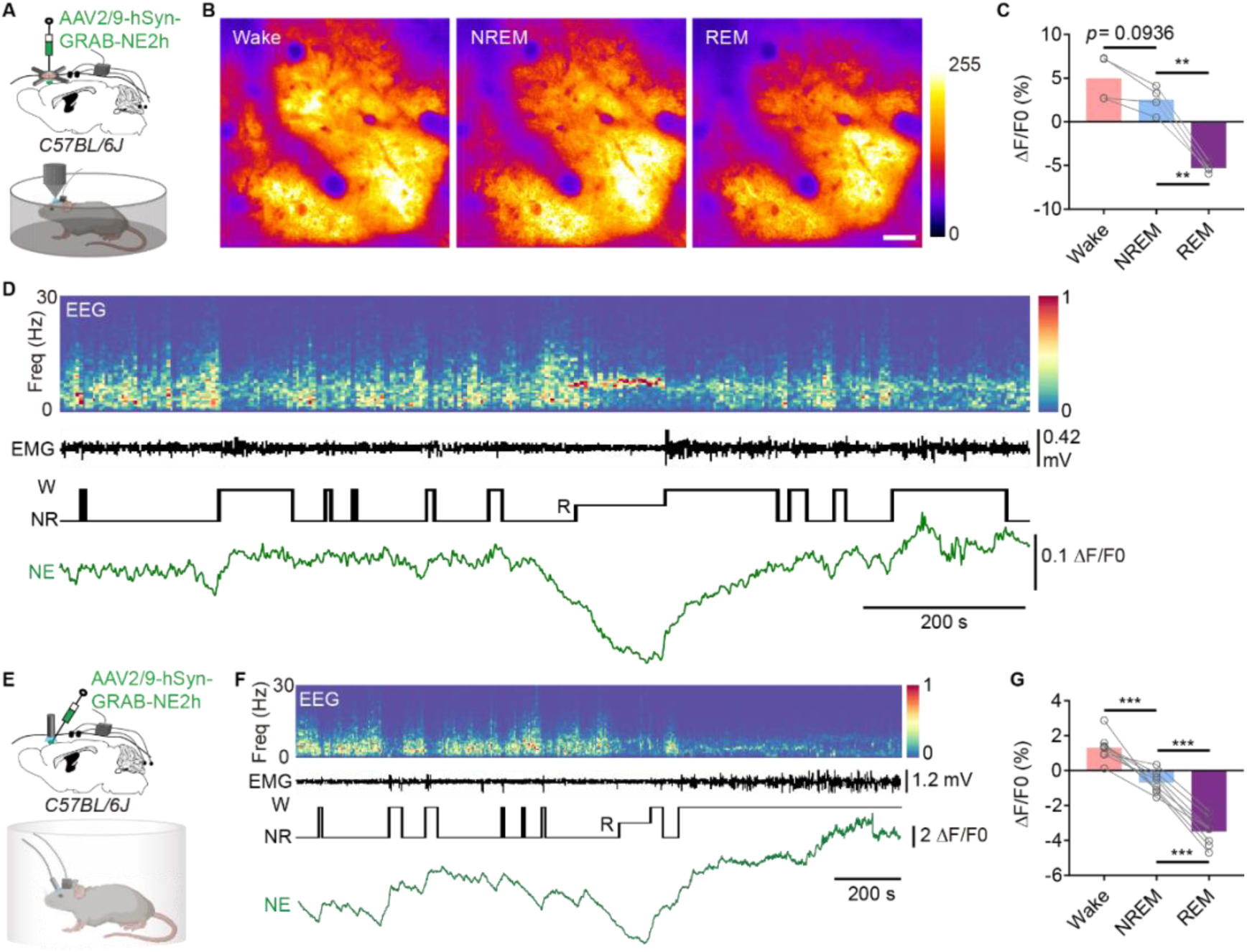
Cortical norepinephrine levels increase during wakefulness. (A) Schematic of the experimental design for examining NE signal during the sleep-wake cycle via head-fixed two-photon imaging. (B) Representative two-photon images of cortical NE biosensor fluorescence during wake, NREM sleep, and REM sleep. Scale bar, 50 μm. (C) Quantification of NE sensor fluorescence (ΔF/F0) across sleep stages (n = 4 mice). (D) Representative EEG power spectrum, EMG trace, hypnogram, and relative changes in NE sensor fluorescence (ΔF/F0) across sleep stages from a single mouse. (E) Schematic of the experimental procedure for fiber photometry combined with EEG–EMG recording to examine NE signal during the sleep-wake cycle. (F) Representative EEG power spectrum, EMG trace, hypnogram, and relative changes in NE signal from fiber photometry (ΔF/F0) across sleep stages from a single mouse. (G) Quantification of NE signal (ΔF/F0) across sleep stages (n = 8 mice). Data are presented as mean ± SEM. Statistical significance was determined using one-way ANOVA followed by Tukey’s post hoc test, *P < 0.05; **P < 0.01; *** P < 0.001; **** P < 0.0001. Each circle indicates an individual mouse.

**Figure S3.**
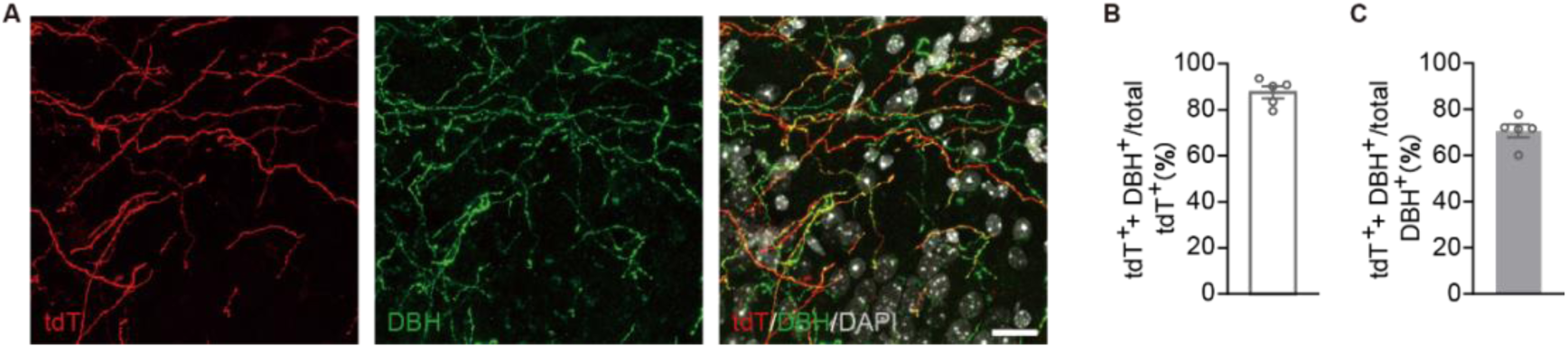
Efficiency of AAV-mediated labeling of cortical NE axons. (A) Representative images showing AAV-labeled NE axons (tdTomato), DBH staining, and the merged image with DAPI. Scale bar, 20 μm. (B) Quantification of the proportion of tdT⁺/DBH⁺ axons relative to total tdT⁺ axons in the cortex (n = 5 mice). (C) Quantification of the proportion of tdT⁺/DBH⁺ axons relative to total DBH⁺ axons in the cortex (n = 5 mice). Data are presented as mean ± SEM.

**Figure S4.**
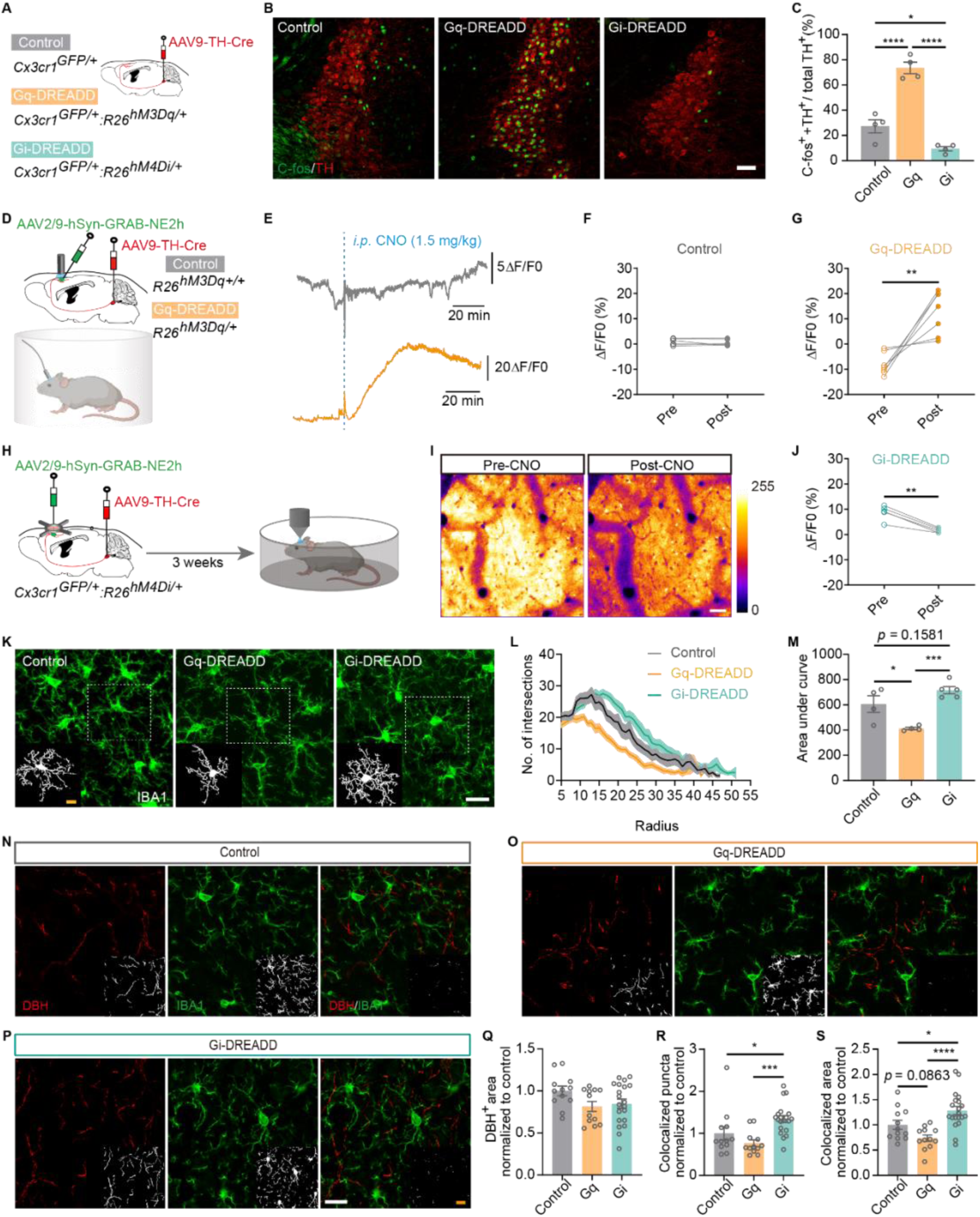
Chemogenetic manipulation of LC-NE neurons controls microglial complexity and their interactions with NE axons. (A) Schematic of transgenic mouse lines and viral injection procedures in the control, Gq-DREADD, and Gi-DREADD groups. (B) Representative immunohistochemical images showing C-fos and TH staining in the LC region from control, Gq-DREADD, and Gi-DREADD groups. Scale bars, 50 μm. (C) Quantification of the proportion of C-fos⁺/TH⁺ neurons relative to total TH⁺ neurons in the LC region across groups (n = 4 mice per group). (D) Schematic of the experimental procedure for fiber photometry recording to assess NE signals under chemogenetic manipulation. (E) Representative NE signal traces (ΔF/F0) before and after CNO intraperitoneal injection in control and Gq-DREADD group mice. (F, G) Quantification of NE signals (ΔF/F0) before and after CNO injection in control (F, n = 6 mice) and Gq-DREADD (G, n = 7 mice) groups. (H) Schematic of the experimental design to examine the NE signal in the Gi-DREADD group via head-fixed two-photon imaging. (I) Representative two-photon images of cortical NE biosensor fluorescence before and after CNO injection. Scale bar, 50 μm. (J) Quantification of NE sensor fluorescence (ΔF/F0) before and after CNO injection (n = 5 mice). (K) Representative immunohistochemical images of cortical IBA1 staining in control, Gq-DREADD, and Gi-DREADD groups (Scale bars, 20 μm). White boxes indicate representative IBA1 mask (Scale bars, 10 μm). (L) The number of intersections per microglial cell, measured on concentric circles at 5 μm intervals from the soma, obtained from Sholl analysis (control, 16 microglia from 4 mice; Gq, 31 microglia from 4 mice; Gi, 31 microglia from 5 mice). (M) Quantification of the area under the curve shown in (L) for the three groups (Control and Gq group, n=4 mice; Gi, n=5 mice). (N-P) Representative immunohistochemical images of DBH and IBA1 staining in the cortex from control, Gq-DREADD, and Gi-DREADD groups. Scale bars, 20 μm. Insets in the lower right corner display DBH, IBA1, and their colocalization masks. Scale bar, 20 μm. (Q) Quantification of DBH⁺ area in control, Gq-DREADD, and Gi-DREADD groups (Control and Gq, n = 12 slices from 4 mice; Gi, n = 16 slices from 5 mice). (R) Quantification of colocalized DBH⁺ and IBA1⁺ puncta in control, Gq-DREADD, and Gi-DREADD groups (Control and Gq, n = 12 slices from 4 mice; Gi, n = 16 slices from 5 mice). (S) Quantification of the colocalized area of DBH⁺ and IBA1⁺ signals in control, Gq-DREADD, and Gi-DREADD groups (Control and Gq, n = 12 slices from 4 mice; Gi, n = 16 slices from 5 mice). Data are presented as mean ± SEM. Statistical significance was determined using one-way ANOVA followed by Tukey’s post hoc test in C, M and S; using one-way ANOVA followed by Dunn’s post hoc test in Q, R; paired t test in F, G and J; *P < 0.05; **P < 0.01; *** P < 0.001; **** P < 0.0001. Each circle indicates an individual mouse in C, F, G, J and M. Each circle indicates an individual slice in Q, R and S.

**Figure S5.**
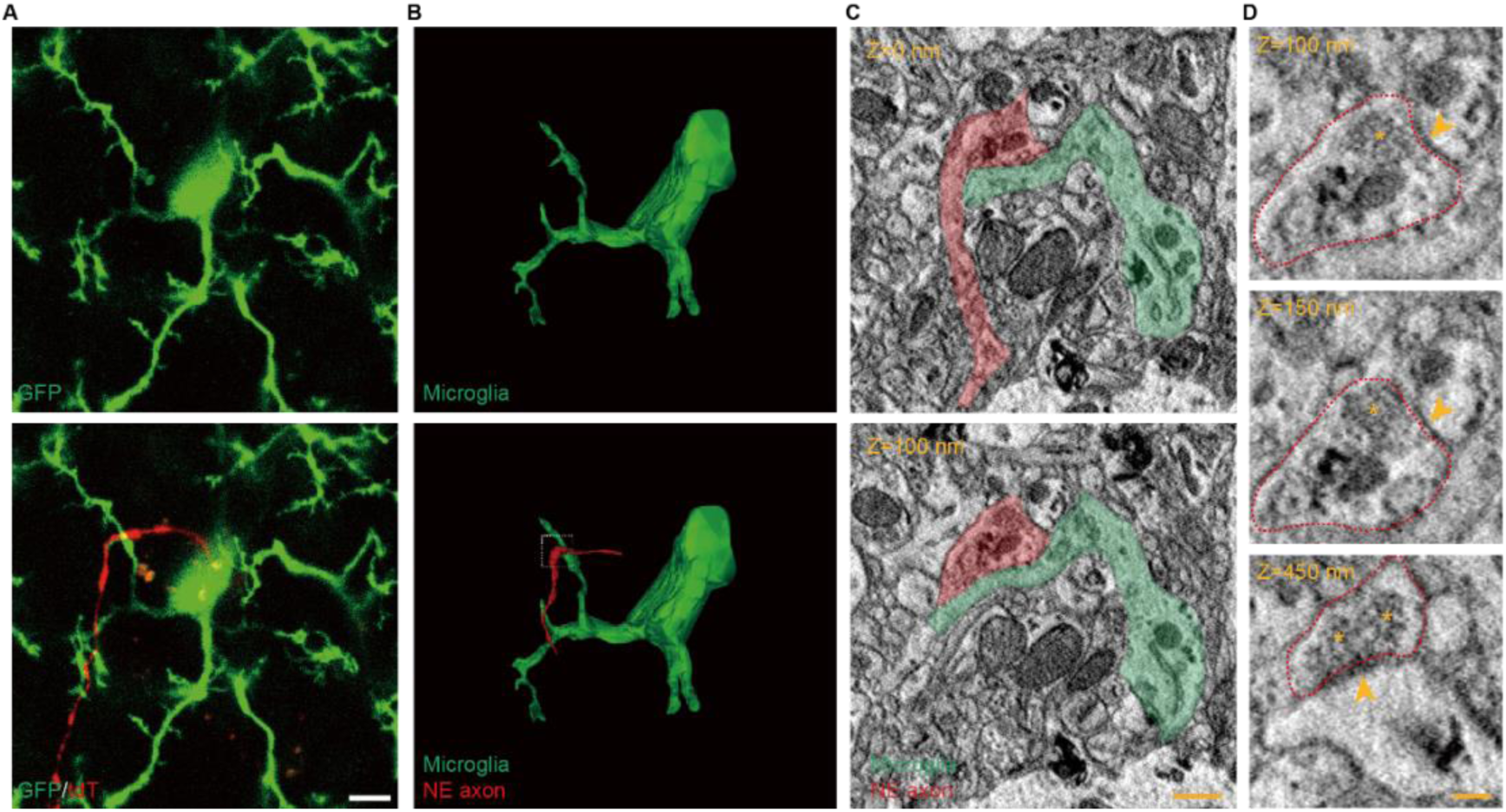
3D electron microscopy reveals that microglial filopodia interact with NE axonal boutons. (A) Representative confocal images showing microglia (GFP) with NE axon (tdTomato) interactions in the cortex, followed by electron microscopy reconstruction. Scale bar, 5 μm. (B) 3D serial reconstruction of a microglia (green) with NE axon (red) interaction. White boxes indicate regions magnified in (C). (C) Electron micrographs showing a microglial process (green) contacting an NE axon (red) at the ultrastructural level using SEM. Scale bar, 500 nm. (D) Electron micrographs showing an NE bouton making synapses with the dendritic spine, as indicated by the arrowheads. Red dashed lines outline the NE bouton, and the yellow asterisk indicates sites of vesicle clustering. Scale bar, 200 nm.

**Figure S6.**
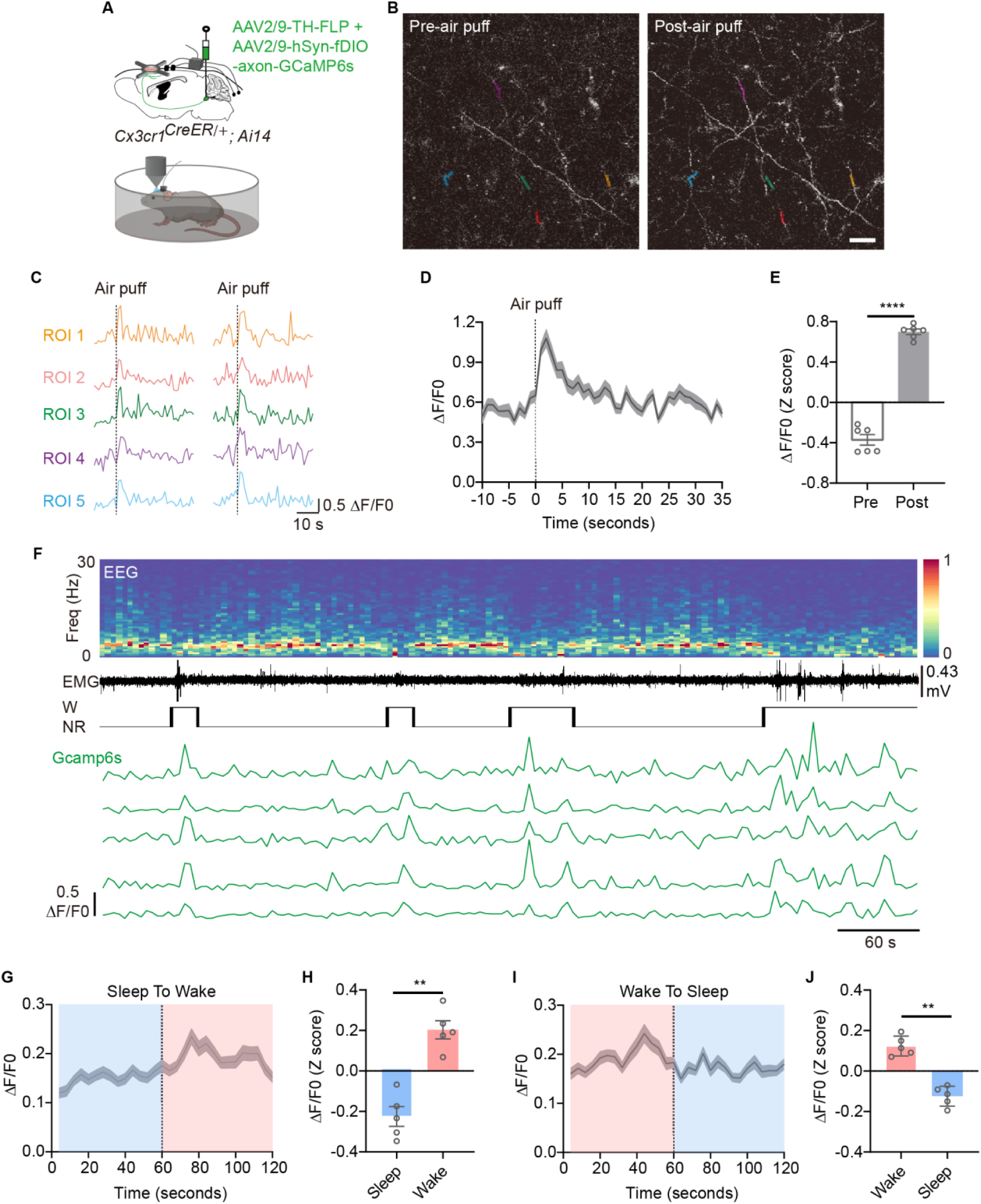
Increased Ca²⁺ activity of NE axons in response to air puff stimulation or during wakefulness. (A) Experimental design for examining Ca²⁺ activity of NE axons in the frontal cortex. (B) Representative two-photon images showing Ca²⁺ signals in NE axons before and after air puff stimulation. Different colored segments correspond to the Ca²⁺ signal traces shown in (C). Scale bar, 20 μm. (C) Representative Ca²⁺ signal traces from five ROIs indicated in (B). (D) Average Ca²⁺ activity (ΔF/F0) across all ROIs of NE axons in response to air puff stimulation (87 ROIs from 6 mice). (E) Quantification of Ca²⁺ activity (ΔF/F0, Z score) before and after air puff stimulation (n=6 mice). (F) Representative EEG power spectrum, EMG trace, hypnogram, and relative Ca²⁺ activity (ΔF/F0) from two-photon imaging across sleep–wake states in a single mouse. (G) Average Ca²⁺ activity (ΔF/F0) across all ROIs of NE axons during the sleep-to-wake transition (78 ROIs from 5 mice). (H) Quantification of Ca²⁺ activity (ΔF/F0) during sleep and wake episodes (n=5 mice). (I) Average Ca²⁺ activity (ΔF/F0) across all ROIs of NE axons during the wake-to-sleep transition (86 ROIs from 5 mice). (J) Quantification of Ca²⁺ activity (ΔF/F0) during wake and sleep episodes (n=5 mice). Data are presented as mean ± SEM. Statistical significance was determined using a paired t-test in all graphs, *P < 0.05; **P < 0.01; *** P < 0.001; **** P < 0.0001. Each circle indicates an individual mouse.

**Figure S7.**
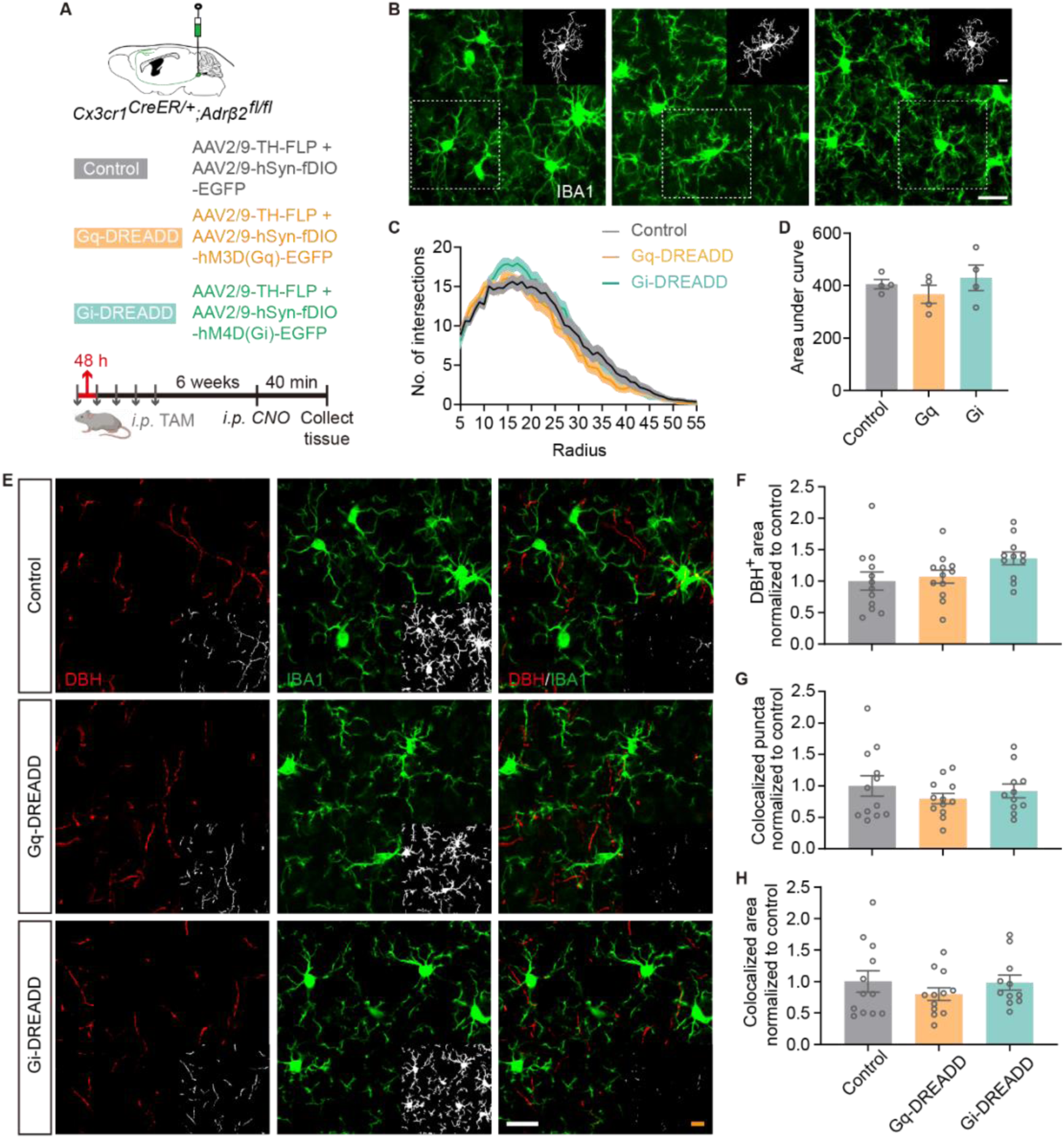
Microglial β2AR deficiency abolishes changes in microglial complexity and microglia–NE axon interactions following chemogenetic manipulation of LC-NE neurons. (A) Schematic of viral injection procedures in control, Gq-DREADD, and Gi-DREADD groups in β2AR cKO mice, and timeline of tamoxifen administration. (B) Representative immunohistochemical images of cortical IBA1 staining in control, Gq-DREADD, and Gi-DREADD groups (Scale bars, 20 μm). White boxes indicate representative IBA1 mask (Scale bars, 10 μm). (C) The number of intersections per microglial cell, measured on concentric circles at 5 μm intervals from the soma, obtained from Sholl analysis (35 microglia from 4 mice in each group). (D) Quantification of the area under the curve shown in (C) for the three groups (Control and n=4 mice for each group). (E) Representative immunohistochemical images of DBH and IBA1 staining in the cortex from control, Gq-DREADD, and Gi-DREADD groups. Scale bars, 20 μm. Insets in the lower right corner display DBH, IBA1, and their colocalization masks. Scale bar, 20 μm. (F) Quantification of DBH⁺ area in control, Gq-DREADD, and Gi-DREADD groups (Control and Gq, n = 12 slices from 4 mice; Gi, n = 11 slices from 4 mice). (G) Quantification of colocalized DBH⁺ and IBA1⁺ puncta in control, Gq-DREADD, and Gi-DREADD groups (Control and Gq, n = 12 slices from 4 mice; Gi, n = 11 slices from 4 mice). (H) Quantification of the colocalized area of DBH⁺ and IBA1⁺ signals in control, Gq-DREADD, and Gi-DREADD groups (Control and Gq, n = 12 slices from 4 mice; Gi, n = 11 slices from 4 mice). Data are presented as mean ± SEM. Statistical significance was determined using one-way ANOVA followed by Tukey’s post hoc test in D, F, G and H, *P < 0.05; **P < 0.01; *** P < 0.001; **** P < 0.0001. Each circle represents one brain slice.

**Figure S8.**
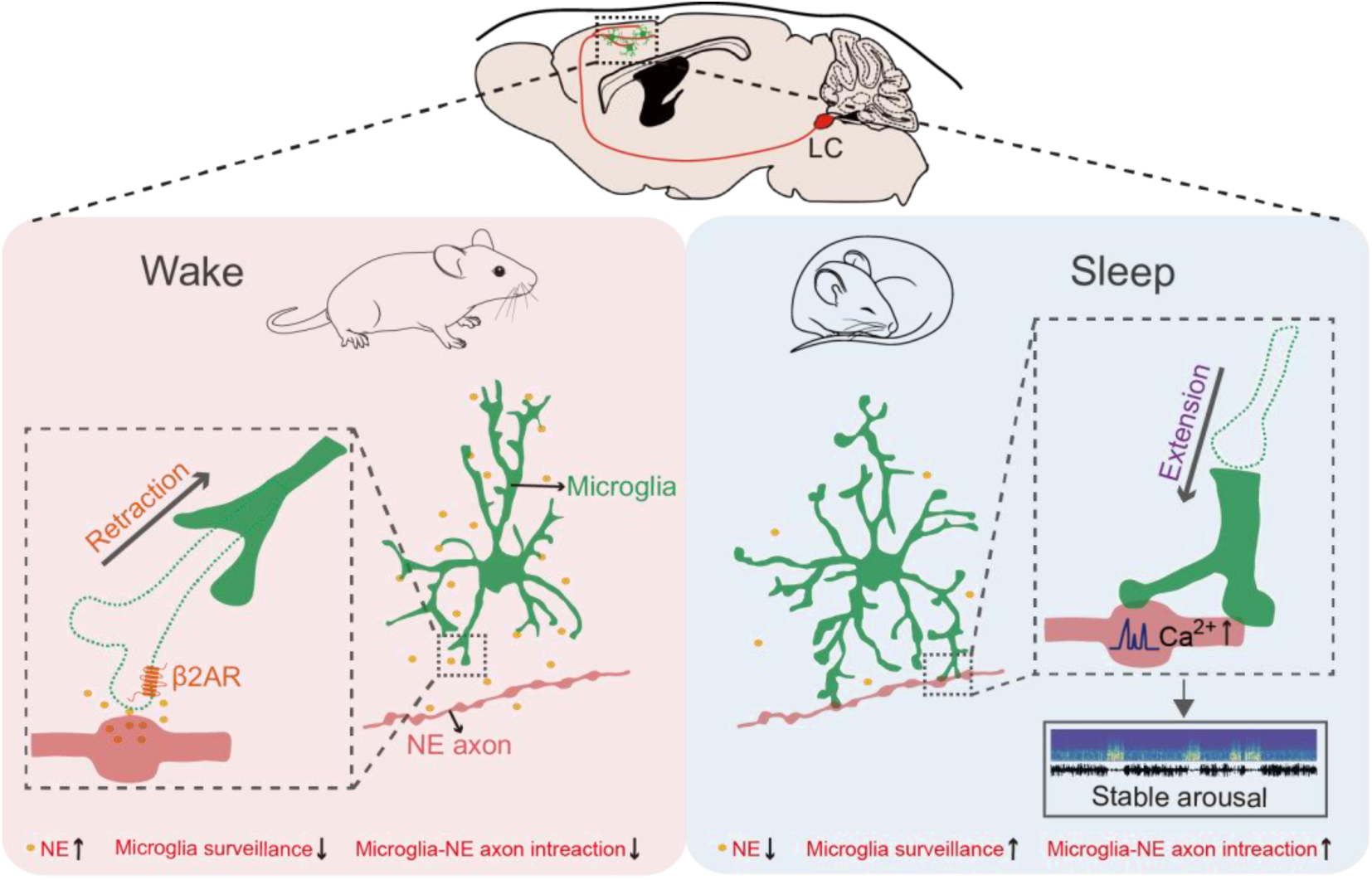
Graphic summary. During sleep, microglia extend their processes in response to reduced NE levels in the brain parenchyma, and their bulbous endings interact with NE axons. These bulbous endings primarily contact NE boutons, enhancing local Ca²⁺ signals in NE axons and contributing to the maintenance of wakefulness.

## Supplementary Videos

**Video S1**. Time-lapse two-photon imaging showing changes in microglial (GFP) process dynamics across wakefulness and sleep states. Related to Figure 1.

**Video S2**. Two-photon imaging of microglia–NE axon (microglia labeled by GFP, NE axons labeled by tdTomato) interactions during wake and sleep. Arrowheads mark newly established contact sites with NE boutons during sleep. Related to Figure 3.

**Video S3**. Two-photon imaging showing Ca²⁺ signaling in NE axons of the frontal cortex. Related to Figure S7.

**Video S4**. Two-photon imaging showing that microglial (tdTomato) contact enhances Ca²⁺ signaling (GFP) in NE axons. Arrowheads indicate contact sites. Related to Figure 5.

## Notes

### Competing Interest Statement

The authors have declared no competing interest.

